# Structure and activation of the human autophagy-initiating ULK1C:PI3KC3-C1 supercomplex

**DOI:** 10.1101/2023.06.01.543278

**Authors:** Minghao Chen, Thanh N. Nguyen, Xuefeng Ren, Grace Khuu, Annan S. I. Cook, Yuanchang Zhao, Ahmet Yildiz, Michael Lazarou, James H. Hurley

## Abstract

The unc-51-like kinase protein kinase complex (ULK1C) is the most upstream and central player in the initiation of macroautophagy in mammals. Here, the cryo-EM structure of the human ULK1C core was determined at amino acid residue-level resolution. A moderate resolution structure of the ULK1C core in complex with another autophagy core complex, the class III phosphatidylinositol 3-kinase complex I (PI3KC3-C1) was also determined. The two complexes co-assemble through extensive contacts between the FIP200 scaffold subunit of ULK1C and the VPS15, ATG14, and BECN1 subunits of PI3KC3-C1.

The FIP200:ATG13:ULK1 core of ULK1C undergoes a rearrangement from 2:1:1 to 2:2:2 stoichiometry in the presence of PI3KC3-C1. This suggests a structural mechanism for the initiation of autophagy through formation of a ULK1C:PI3KC3-C1 supercomplex and dimerization of ULK1 on the FIP200 scaffold.

## Introduction

Macroautophagy (hereafter, “autophagy”) is the main cellular mechanism for the disposal of molecular aggregates and damaged or unneeded organelles ^1^. While autophagy was first characterized as a bulk response to starvation, it is now clear that many forms of bulk and selective autophagy are central in development and cellular homeostasis ^1^. The homeostatic necessity of autophagy is clearest in neurons, which are post-mitotic and so uniquely susceptible to toxic aggregates and damaged organelles ^2^. Autophagic dysfunction has been linked to all major neurodegenerative diseases ^2^, with the clearest genetic linkage to Parkinson’s disease ^3^.

All forms of canonical autophagy, bulk and selective, are initiated upon the recruitment and activation of the FIP200 protein ^4^ and the class III phosphatidylinositol 3-kinase complex I (PI3KC3-C1) ^5–7^. FIP200 serves as the central scaffolding subunit of the ULK1C, providing a platform for the other three subunits: the ULK1 kinase, ATG13, and ATG101 ^8–12^. PI3KC3-C1 contains one copy each of the lipid kinase VPS34, the pseudokinase VPS15, and the regulatory subunits BECN1 and ATG14 ^5,13^. The former three subunits are also present in the PI3K complex II (PI3KC3-C2) involved in endosomal sorting and late steps in autophagy, while ATG14 is uniquely involved in autophagy initiation ^5–7^.

Atomic and near-atomic resolution structures are known for various fragments of these complexes ^14–16^ and low to moderate resolution structures are known for human PI3KC3-C1 ^17,18^ and the core of human ULK1C ^19^.

The means by which ULK1C and PI3KC3-C1 activities are switched on and off are critically important for the physiology of autophagy initiation and for therapeutic interventions targeting neurodegeneration ^20^. Yet these mechanisms have thus far been hidden because of the limitations of the available fragmentary or low-resolution structures. While ULK1C and PI3KC3-C1 are both critical for autophagy initiation and are both recruited at the earliest stages, beyond the presence of ULK1 phosphorylation sites on PI3KC3-C1 subunits ^21,22, 23,24^, it has been unclear how their activities are coordinated. Here we report the structures of the human ULK1C core at resolutions adequate for amino acid residue-level interpretation. We also show that ULK1C and PI3KC3-C1 form a physical supercomplex, determine its structure, and show that ULK1C can undergo a stoichiometric switch leading to the recruitment of two copies of the ULK1 kinase itself.

## Results

### Cryo-EM structure of a 2:1:1 stoichiometric ULK1C core

The ordered core of ULK1C was purified in a form consisting of the FIP200 N-terminal domain (1-640) and the ULK1 C-terminal microtubule-interacting and transport (MIT) domain (836-1059) fused with the ATG13 residues (363-517) responsible for binding to FIP200 and ULK1 ^19^ (Figure 1A). The peripheral domains, including the coiled-coil domain (791-1498) and the claw domain (1499-1594) of FIP200, the kinase domain of ULK1 (1-278) and the ATG13 (1-190):ATG101(1-218) HORMA domain, were removed from the construct because they are flexibly connected to the ULK1C core only via disordered loops and not known to participate in the ordered core assembly. Deletion of these domains does not affect the stability of the ULK1C core and facilitated its structure determination. A monodisperse peak from size exchange chromatography (SEC) was collected and used for cryo-EM data collection. Image processing and 3D reconstruction resulted in a cryo-EM density map with a local resolution of 3.35 Å in the best regions, including the FIP200:ATG13:ULK1 interface (Figure. S1, A to G). The distal tips of the FIP200^NTD^ molecules are mobile and the density there is less defined. The ATG13:ULK1 unit resembles the yeast Atg13^MIM^:Atg1^MIT^ complex ^25^, so we adopted the MIT/MIM terminology for the human complex. The quality of the density allowed assignment of amino acid residues of the ULK1^MIT^ domain and ATG13^MIM^ (Figure. S1, H to L). The structure confirmed the previous observation of a 2:1:1 FIP200:ATG13:ULK1 complex ^19^, while defining in residue-level detail how ULK1^MIT^ and ATG13^MIM^ bind to FIP200.

**Figure 1.**
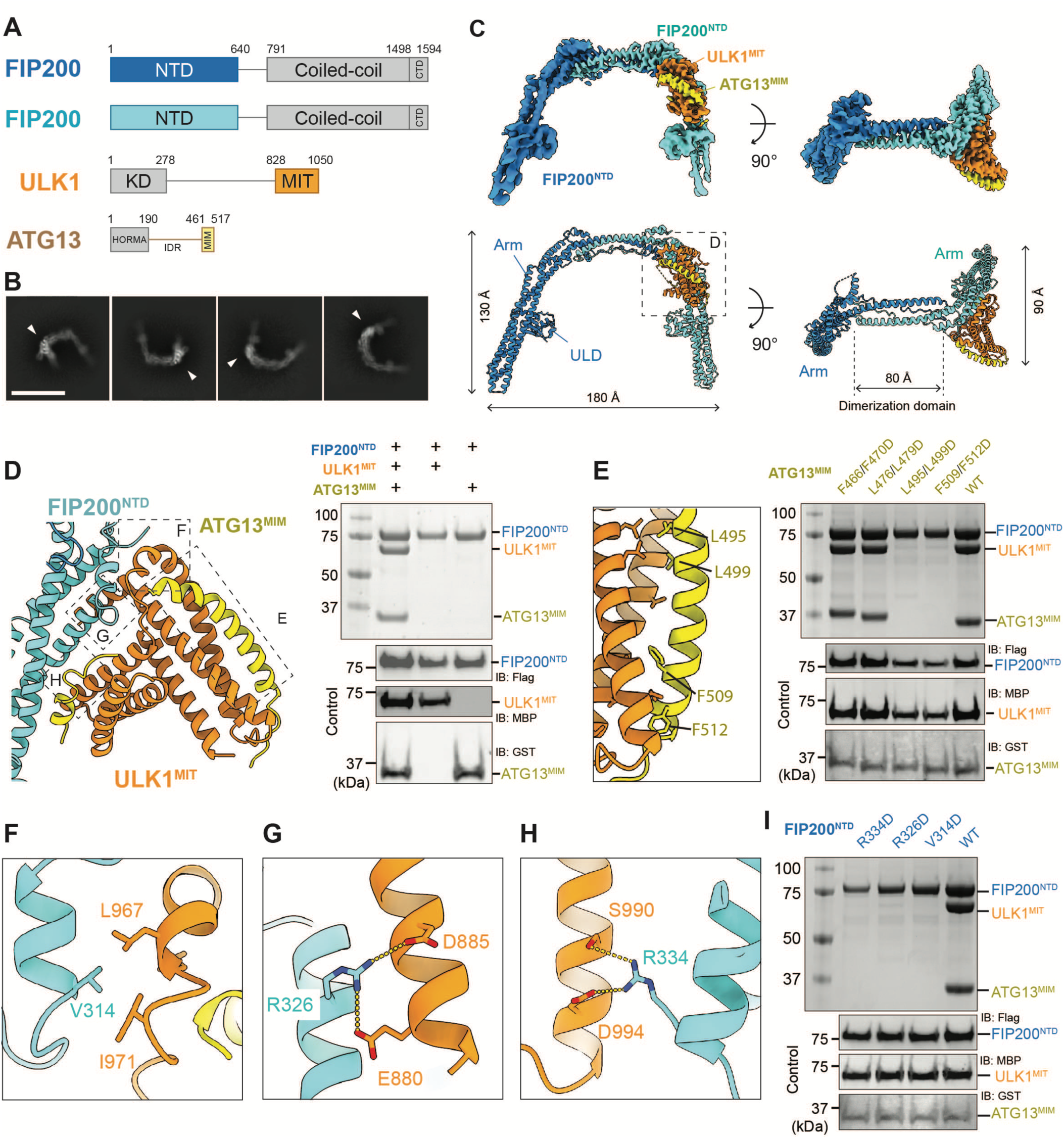
Structure of the ULK1C core. **(A)** Domain organization of the ULK1C core. Gray indicates the regions truncated in this study. NTD: N-terminal domain, CTD: C-terminal domain, KD: Kinase domain, MIT: Microtubule-interacting and transport domain, HORMA: Hop1p/Rev7p/MAD2 domain, MIM: MIT-interacting motif. (**B**) Representative reference-free 2D class averages of the ULK1C core. Scale bar: 20 nm. (**C**) Overview of the EM map (upper) and the coordinates (lower) of the ULK1C core. Subunits are depicted in the same color as in (**A**). The map is contoured at 12σ. (**D**) Pull-down assay of FIP200^NTD^ with ULK1^MIT^ and ATG13^MIM^. Strep-Tactin resin was loaded with TSF (TwinStrep-Flag)-tagged FIP200^NTD^ to pull-down MBP-tagged ULK1^MIT^ and GST-tagged ATG13^MIM^ in various combinations as indicated above the lanes. (**E**) Close-up view of the interface between ULK1^MIT^ and ATG13^MIM^ and the corresponding Strep pull-down assay. FIP200^NTD^-TSF was used to pull-down various GST-tagged ATG13 mutants (F466D/F470D, L476D/L479D, L495D/L499D, and F509D/F512D) and MBP-tagged wild type ULK1^MIT^. (**F-H**) Close-up views of the interface between ULK1^MIT^ and FIP200. Key residues for the binding interface are indicated, with hydrogen bonds shown as yellow dotted lines. (**I**) Strep pull-down assay for the FIP200-ULK1 interface. Various FIP200^NTD^-TSF mutants (R334D, R326D, and V314D) were used to pull-down the wild type ULK1^MIT^ and ATG13^MIM^. All pull-down results were visualized by SDS-PAGE and Coomassie blue staining.

The ULK1C core has dimensions of 180 × 130 × 90 Å, contains two molecules of FIP200^NTD^ and one molecule each of ULK1 and ATG13 (Figure 1B and 1C). FIP200 ^NTD^ forms a C-shaped dimer as seen at low resolution ^19^, which can now be described in detail with sequence assigned. The two arms of FIP200 consist of 120 Å-long bundles formed by three twisted helixes (81-495). The two arms are connected by an 80 Å-long dimerization domain (496-599) bent at nearly 90° to the arms, resulting in the C-shape. The structure after residue 599 was not resolved due to presumed disorder. The FIP200 dimer is in some ways reminiscent of the structure of its yeast counterpart, Atg17 ^26^, although the protein folds are distinct. Atg17 also dimerizes via two arms that are in the same plane, although the arms are arranged in an S-shape instead of a C (Figure S2A). Residues 1-80 of FIP200 form a ubiquitin-like domain (ULD) located at the middle of the Arm domain in the inner side of the C-shape. The higher resolution analysis here confirms the previous observation that the ULD and the Arm domain of FIP200 have the same fold as the scaffold domain of the Tank-binding kinase 1 (TBK1) ^19^ (Figure S2B), which is itself central to the initiation of some forms of autophagy ^27^.

Cryo-EM density for one copy of a ULK1^MIT^:ATG13^MIM^ heterodimer was observed on one “shoulder” of the FIP200 dimer. The ATG13^MIM^ binds collaboratively to both FIP200 and ULK1^MIT^ (Figure 1D), consistent with the role of ATG13 in recruiting ULK1 ^28^ downstream of FIP200. As seen for yeast Atg13:Atg1 ^25^, the structure consists of two four-helix bundles (Figure S2C). Three helices of each are from the ULK1^MIT^ and one from the ATG13^MIM^. Four pairs of hydrophobic residues from the ATG13^MIM^ form prominent direct interactions with ULK1^MIT^ (Figure 1E). The ULK1^MIT^:ATG13^MIM^ heterodimer binds to FIP200 via two main interfaces (Figure 1F to 1H). Three FIP200 residues, Val314, Arg326, and Arg334, were identified by cryo-EM as potentially important for the ULK1:ATG13 interaction, which was confirmed by mutagenesis and pull-down assays (Figure 1I).

### The ATG13^IDR:^FIP200^NTD^ interface

The C-terminal portion of the ATG13 intrinsically disordered region (ATG13^IDR^), corresponding to residues 363-460, was previously demonstrated to bind to the FIP200^NTD 19^, but was not visualized with sufficient continuity to assign amino acid sequence on the basis of the density alone. In this study, we utilized AlphaFold2 ^29^ as corroborated by hydrogen-deuterium exchange mass spectrometry (HDX-MS)^19^ to map four potential binding sites on ATG13 (site 1(370-387), site 2(392-408), site 3(409-414)), and site 4(450-460)) (Figure 2A, 2B and S3). The sites augment the role of the ATG13^MIM^ motif (460-517), referred to as site M hereafter. The interaction of a single ATG13^IDR^ extends through both protomers of the FIP200^NTD^ dimer. Sites 1 and 4 are unique in ATG13, but their binding sites on two FIP200 protomers are structurally symmetric. One FIP200 molecule is occupied by ATG13 site 4 and the ULK1^MIT^:ATG13^MIM^ heterodimer, and the other FIP200 molecule is occupied by site 1-3 of the ATG13^IDR^ loop (Figure 2B). This explains the unusual 2:1:1 stoichiometry of the ULK1C by showing in atomistic detail how the FIP200^NTD^ dimer binds a single copy of ATG13 in the complex.

**Figure 2.**
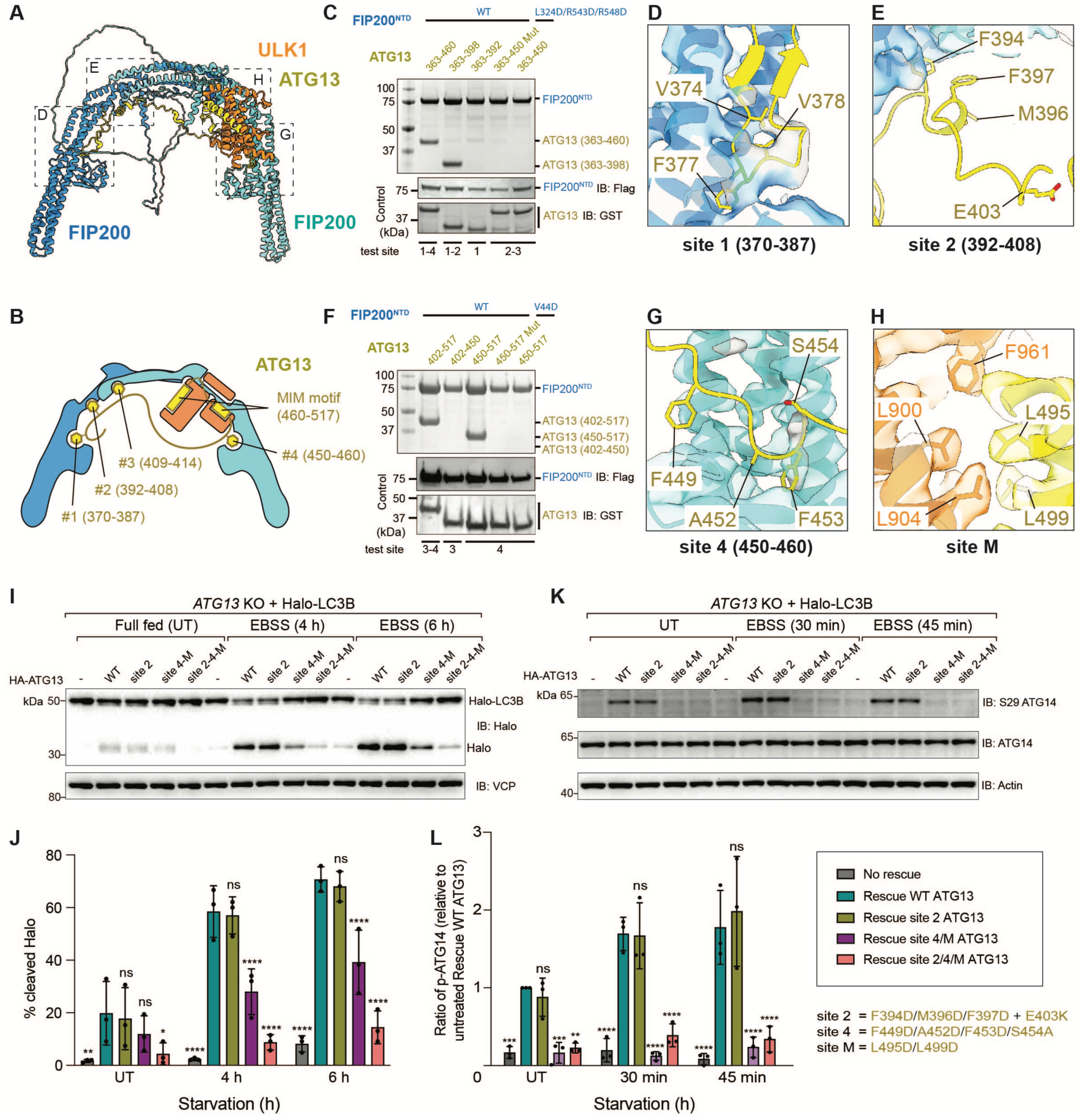
Binding sites of ATG13^IDR^ on FIP200^NTD^. (**A**) AlphaFold2 prediction model of the ULK1C core. (**B**) Schematic diagram of the ULK1C core. (**C**) Truncation and mutagenesis screening conducted by pull-down assay. Strep-Tactin resin was loaded with wild type FIP200^NTD^-TSF to pull-down various ATG13 fragments (363-460 (includes site 1-4), 363-398 (site 1-2), 363-392 (site 1), and 363-450 with F394D/F397D/E403K mutations (mutated site 2)), or loaded with FIP200^NTD^-TSF variant (L324D/R543D/R548D) to pull-down wild type ATG13 fragment 363-450. (**D** and **E**) Close up views of the binding site 1 and 2 of the AlphaFold2 model with the key binding residues shown. The map is contoured at 11σ. (**F)** Strep pull-down assay. The wild type FIP200^NTD^-TSF was used to pull-down various ATG13 fragments (402-517 (includes site 3-4), 402-450 (site 3), 450-517 (site 4), and 450-517 with A452D/F453D mutations (mutated site 4)), or FIP200^NTD^-TSF variant (V44D) was used to pull-down wild type ATG13 fragment 450-517 (site 4). The results from the pull-down assays (**C** and **F**) were visualized by SDS-PAGE and Coomassie blue staining. (**G)** Close up view of the binding site 4 of the AlphaFold2 model. (**H**) Close up view of the ULK1^MIT^ binding site (site M) of ATG13^MIM^ determined by cryo-EM structure. The key binding residues are shown. (**I** and **J**) *ATG13* KO cells stably expressing Halo-LC3B without rescue or rescued with different versions of HA-ATG13 were treated with 50 nM TMR-conjugated Halo ligand for 15 min. Following that, cells were washed with 1x PBS and treated with EBSS for indicated time periods, harvested, and analysed by immunoblotting with indicated antibodies (**I**) and the percentage of the cleaved Halo band was quantified (**J**). Data in (**J**) are mean ± SD from three independent experiments. ns = not significant, ****P<0.0001 (two-way ANOVA). The mutation sites of each variant and their corresponding residues are shown in the insert. (**K**) *ATG13* KO cells expressing Halo-LC3B without rescue or rescued with indicated versions of HA-ATG13 were left untreated or treated with EBSS for 30 min or 45 min. Cells were harvested and analysed by immunoblotting with indicated antibodies. (**L**) The ratio of phosphorylated ATG14 (p-ATG14) relative to untreated rescue WT ATG13 samples was quantified. Data in (**L**) are mean ± SD from three independent experiments. ns = not significant, ****P<0.0001 (two-way ANOVA). The definition of mutants, shown at the bottom right, applies to panels **I**-**L**.

We screened a series of truncations to map the FIP200 binding determinants of ATG13^IDR^. Two ATG13 fragments, 363-398 (corresponding to site 2), and 450-460 (site 4) were independently sufficient for FIP200 binding (Figure 2C), consistent with the previous observation that these regions are protected from HDX-MS ^19^ (Figure S3B). Point mutagenesis showed that ATG13 residues Phe394, Phe397, and Glu403, and FIP200 residues Leu324, Arg543, and Arg548, are required for binding *via* site 2 (Figure 2C to 2E), and ATG13 residues Ala452 and Phe453, and FIP200 Val44 are key for the binding site 4 (Figure 2F and 2G). We tested whether disrupting the binding between ATG13 and FIP200 would affect bulk autophagy in response to starvation. ATG13 variants (Figure 2D, 2E, 2G and 2H) were introduced into *ATG13* KO HeLa cells (Figure S4). Mutation of site 2 (F394D/M396D/F397D/E403K) alone did not have a significant effect. Mutation of sites 4 and M (site 4: F449D/A452D/F453D/S454A, site M: L495D/L499D) led to a substantial reduction in flux, while mutations of 2, 4, and M together nearly eliminated flux (Figure 2I and 2J). To investigate whether the reduction in autophagy is due to the decreased activity of ULK1C, we measured the phosphorylation level of ATG14^S29^, a substrate of ULK1 kinase that plays an important role in regulating the function of PI3KC3-C1 ^23^. The results of the phosphorylation assay showed that reduced ULK1 activity correlates with reductions in autophagy flux. Mutation of site 2 alone had no significant effect, while the combined mutations of sites 2, 4 and M eliminated ULK1 activity (Figure 2K and 2L).

### ULK1C and PI3KC3-C1 form a supercomplex

PI3KC3-C1 is a substrate of ULK1 ^21,23^. However, it has been unclear whether PI3KC3-C1 is recruited coordinately with ULK1C in autophagy initiation. To determine if there was a direct interaction, and if so, with which portion of the complex, we assessed binding of PI3KC3-C1 to the ULK1C core and the ATG13-ATG101 HORMA dimer. Direct binding between FIP200 and PI3KC3-C1 was observed by bead binding assay (Figure 3A and 3B). The ATG13-ATG101 HORMA dimer did not interact with PI3KC3-C1 under these conditions (Figure 3A and 3B). ULK1C and PI3KC3-C1 were mixed and visualized by cryo-EM (Figure S5A). A small population of ULK1C core in complex with PI3KC3-C1 was observed in this dataset (Figure S5B). The mixture was then further purified by pull-down with PI3KC3-C1 (Figure 3C) and subjected to cryo-EM (Figure 4 and S6). 2D classes of the ULK1C:PI3KC3-C1 supercomplex were visualized, in which the distinctive C-shaped density of the ULK1C core and the V-shaped density of the PI3KC3-C1 were both clearly observed (Figure 4A).

**Figure 3.**
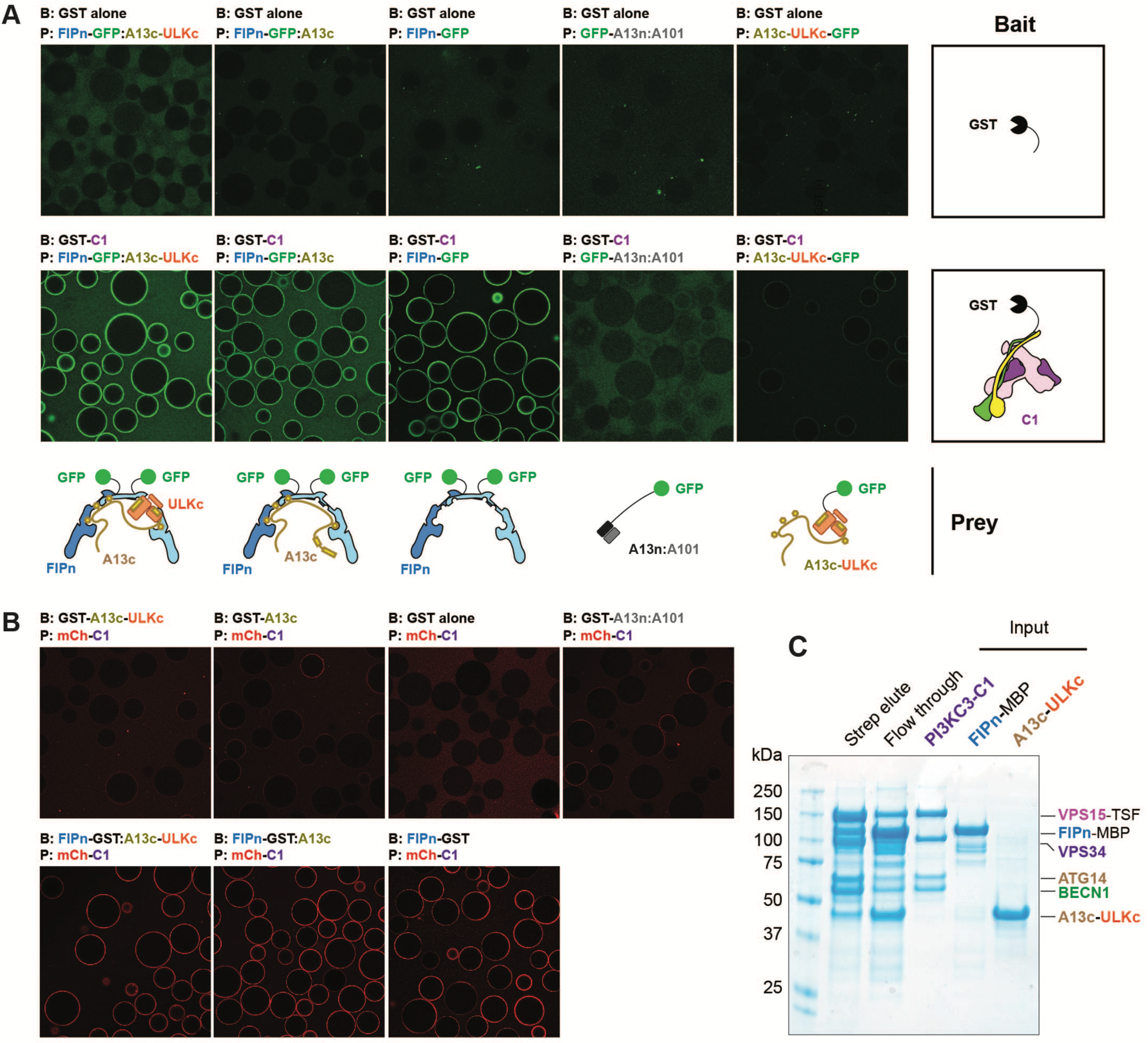
Direct interaction between ULK1C and PI3KC3-C1. (**A**) Beads binding assay of ULK1C core and PI3KC3-C1. GSH beads coated with GST-tag alone or GST-tagged PI3KC3-C1 was used to recruit various combination of GFP-tagged proteins. The schematic drawing illustrates the bait and prey samples used for each condition. (**B**) Complimentary beads binding assay for confirming the interaction between FIP200 and PI3KC3-C1. GST-tagged various proteins were used to recruit the mCherry-tagged PI3KC3-C1. (**C**) SDS-PAGE of the ULK1C: PI3KC3-C1 pull down sample. Strep resin was used to pull down the TSF-tagged VPS15. The eluate was obtained by washing the resin with buffer containing 10 mM d-Desthiobiotin and visualized by SDS-PAGE and Coomassie blue staining.

**Figure 4.**
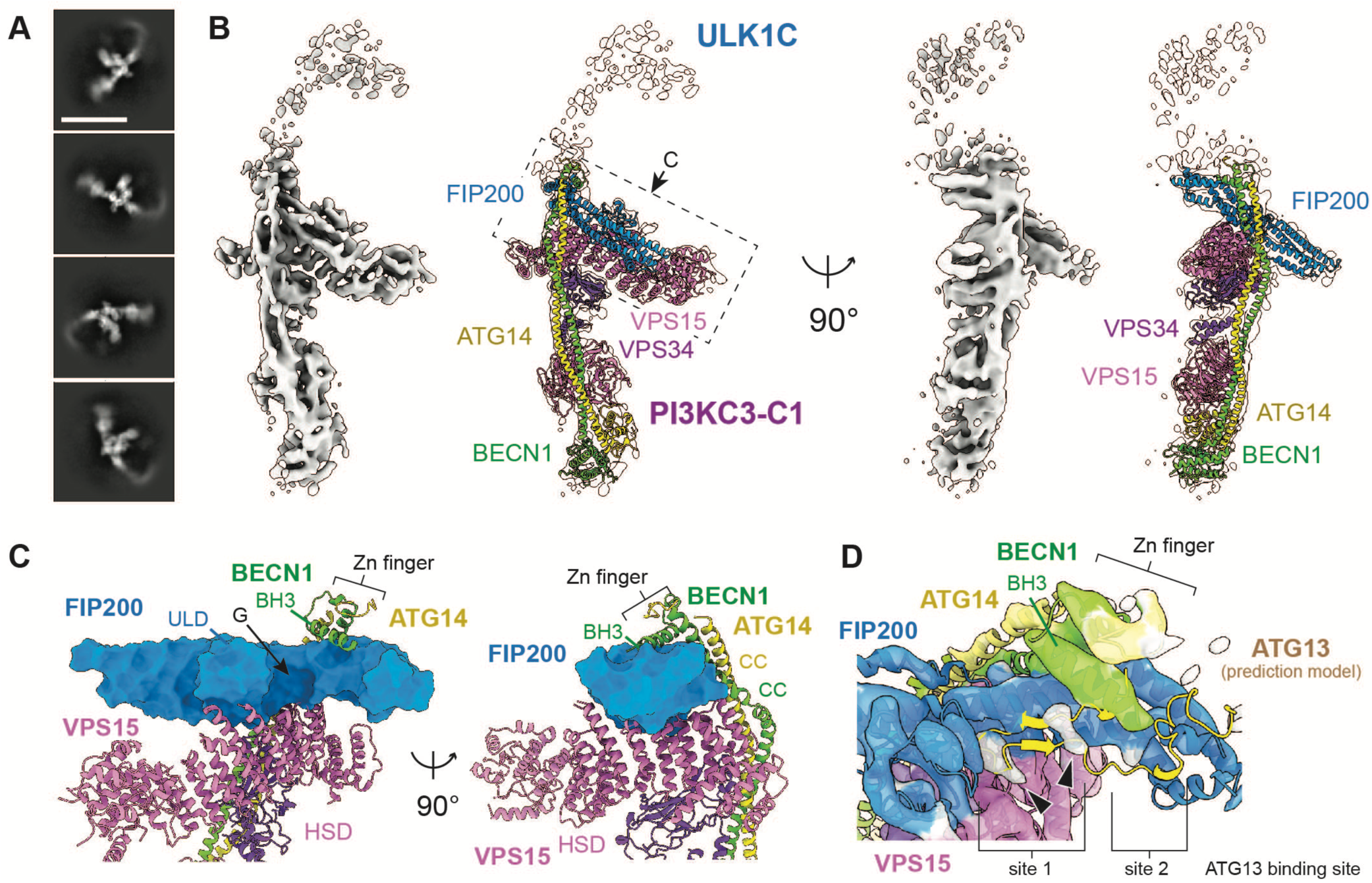
Structure of the ULK1C:PI3KC3-C1 supercomplex. (**A**) Representative 2D class average of the ULK1C:PI3KC3-C1 supercomplex. Scale bar: 20 nm. (**B**) Overview of the EM map and the coordinates of ULK1C:PI3KC3-C1 supercomplex superposed with the EM map contoured at 8σ. Note only one FIP200 molecule of the ULK1C was modeled (blue). (**C**) Close up view of the FIP200 binding interface of PI3KC3-C1. The FIP200 is shown in surface representation and the PI3KC3-C1 is shown in ribbon representation. The FIP200^ULD^, VPS15^HSD^, BECN1, CC and Zn-Finger domains formed by BECN1/ATG14 are indicated. (**D**) Close up view of the unassigned EM densities at site 1 and 2. Local refinement map is shown at 22σ. The assigned map is colored based on the molecule (Blue: FIP200, Pink: VPS15, Green: BECN1, Yellow: ATG14). The extra densities observed in site 1 are shown in gray and indicated by arrows.

Further 3D reconstruction resulted in an EM map containing the crescent FIP200 dimer and all the four subunits (VPS34, VPS15, BECN1, and ATG14) of PI3KC3-C1 (Figure S6). The peripheral regions, including the FIP200 molecule binding with the ULK1^MIT^/ATG13^MIM^ and the VPS34^HELCAT^ (Helical and Catalytic domain) are not modeled due to their high mobility. In contrast, the regions containing VPS15^HSD^ (Helical Solenoid Domain), BECN1^CC^/ATG14^CC^ (Coiled-Coil domain), BECN1^BH3^, and the FIP200 arm not bound to ULK1^MIT^/ATG13^MIM^ (hereafter, the “proximal arm”) are the best resolved parts, with local resolution at 6.84 Å (Figure S6H). This enabled unambiguous identification of the secondary structures identified in the atomistic structures of the PI3KC3-C1 core ^30^ and the proximal arm of FIP200 (Figure 4B and S7A).

Two main interfaces are identified between FIP200 and PI3KC3-C1 (Figure 4C). The first interface is formed between the FIP200^(166–181/473–485)^ and BECN1^BH3^, a well-studied Bcl-2/Bcl-XL binding region that regulates autophagy and apoptosis ^31–34^. A zinc-finger motif, composed of BECN1^C137/C140^ and ATG14^C43/C46^, is located behind the BH3 domain and supports the conformation. The BECN1-ATG14 zinc-finger and the BECN1 BH3 region have not been visualized in previous PI3KC3-C1 structures, thus it appears that the presence of the FIP200 proximal arm induces their ordering. The second interface is formed by the globular ULD domain of FIP200 and the curved HSD domain of VPS15 (Figure S7B). The VPS15^HSD^ adjoins FIP200 sites 1 and 2, forming a large pocket with the FIP200 proximal arm and ULD. Fragmented EM density is observed in this pocket (Figure 4D). Superposition of a predicted model of ULK1C core to the supercomplex structure shows that the unassigned density could be contributed by ATG13^371–385^, although the resolution is insufficient for unambiguous assignment. Overall, the structure shows that PI3KC3-C1 and FIP200 interact extensively and intimately. It is striking that the BECN1^BH3^ and ATG14-BECN1 zinc-finger regions become ordered and wrap more than half-way around the FIP200 proximal arm, and also striking that these regions closely approach ATG13 sites 1-2.

### FIP200 scaffolding of ULK1 dimerization

In the same ULK1C:PI3KC3-C1 mixture data set described above, a particle class of ULK1C showed clear 2:2:2 stoichiometry (Figure 5A and S5C). The presence of the ULK1^MIT^:ATG13^MIM^ heterodimer was confirmed on both of the FIP200 shoulders, demonstrating that the stoichiometry of the ULK1 complex was altered in the cryo-EM sample by the presence of PI3KC3-C1.

**Figure 5.**
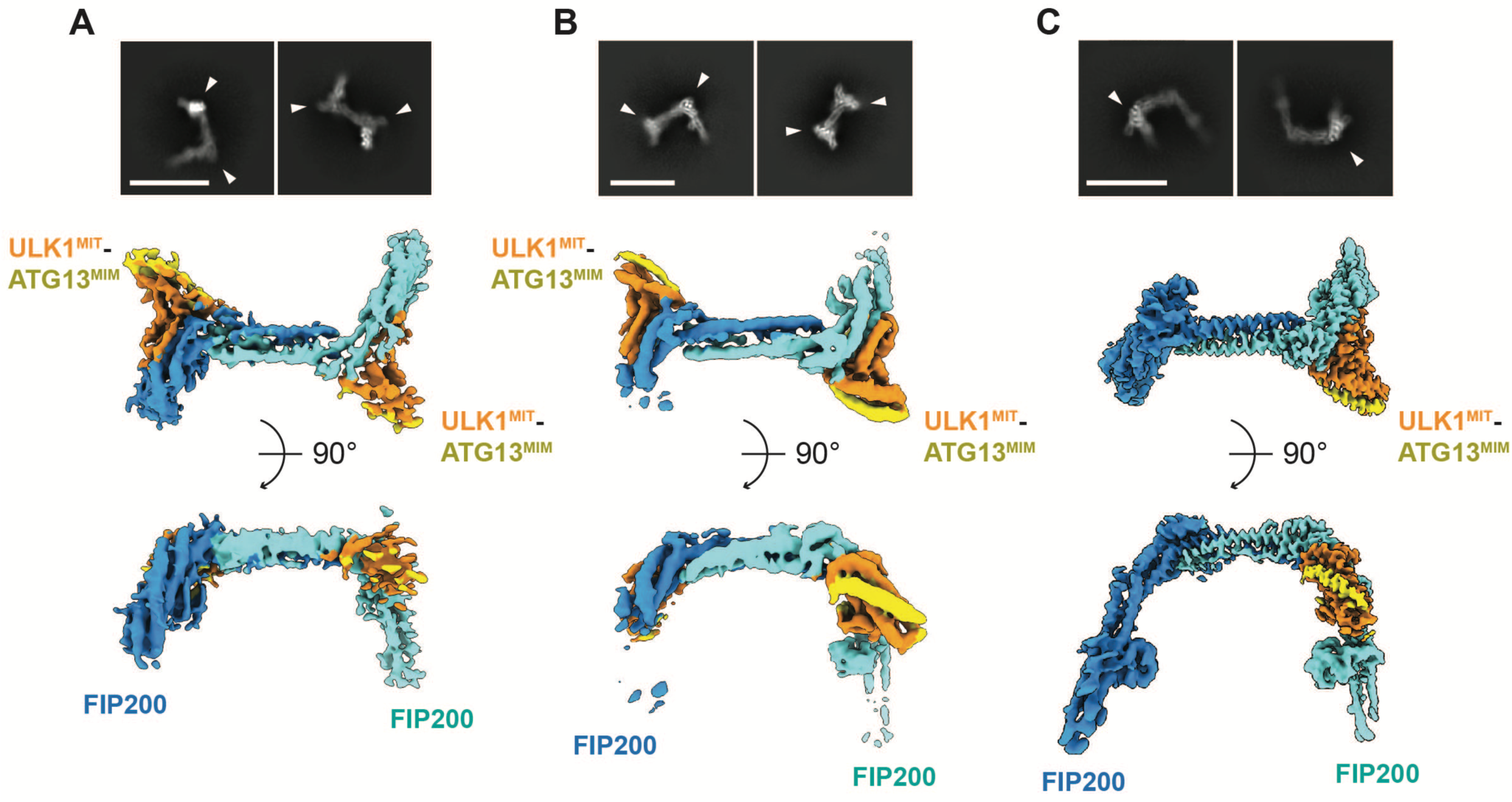
ULK1 dimerization on the FIP200 scaffold. (**A**) Representative 2D class average and the EM map of the ULK1C (2:2:2) core in the presence of PI3KC3-C1. The ULK1^MIT^-ATG13^MIM^ domains are indicated with arrows. Scale bar: 20 nm. The EM map is contoured at 7σ. (**B**) Representative 2D class average and the EM map of the ULK1C (2:2:2) core of the ATG13^450–517^ truncation mutant. The EM map is contoured at 14σ. (**C**) Repeated 2D class averages and the EM map of the ULK1C (2:1:1) core as shown in Figure 1. B and C for comparison. The EM map is contoured at 12σ.

ATG13 site 2 is not involved in the 2:2:2 complex, therefore, we hypothesized that this site might function as a brake to negatively regulate formation of this complex. To test this hypothesis, we truncated ATG13^IDR^ at residue 449 and determined the cryo-EM structure of an ATG13^450–517^-containing version of the ULK1C core in the absence of PI3KC3-C1. The 2D averages clearly show densities ULK1^MIT^:ATG13^MIM^ heterodimer on both sides of the shoulder (Figure 5B), which was confirmed by 3D reconstruction at 4.46 Å (Figure S5, D to H). The stoichiometry was investigated in solution by size-exclusion chromatography with in-line multi-angle light scattering (SEC-MALS) (Figure S8A) and mass photometry (Figure 6). The ULK1C cores in the presence and absence of the ATG13^IDR^ region displayed masses consistent with 2:1:1 and 2:2:2 complexes, respectively, consistent with the cryo-EM data. Furthermore, the ATG13 mutant F394D/F397D/E403K, designed to disrupt site 2, also formed a complex with a 2:2:2 stoichiometry (Figure S8B). This observation demonstrates that site 2 of ATG13^IDR^ opposes the recruitment of the second ULK1 kinase to the FIP200 dimer.

**Figure 6.**
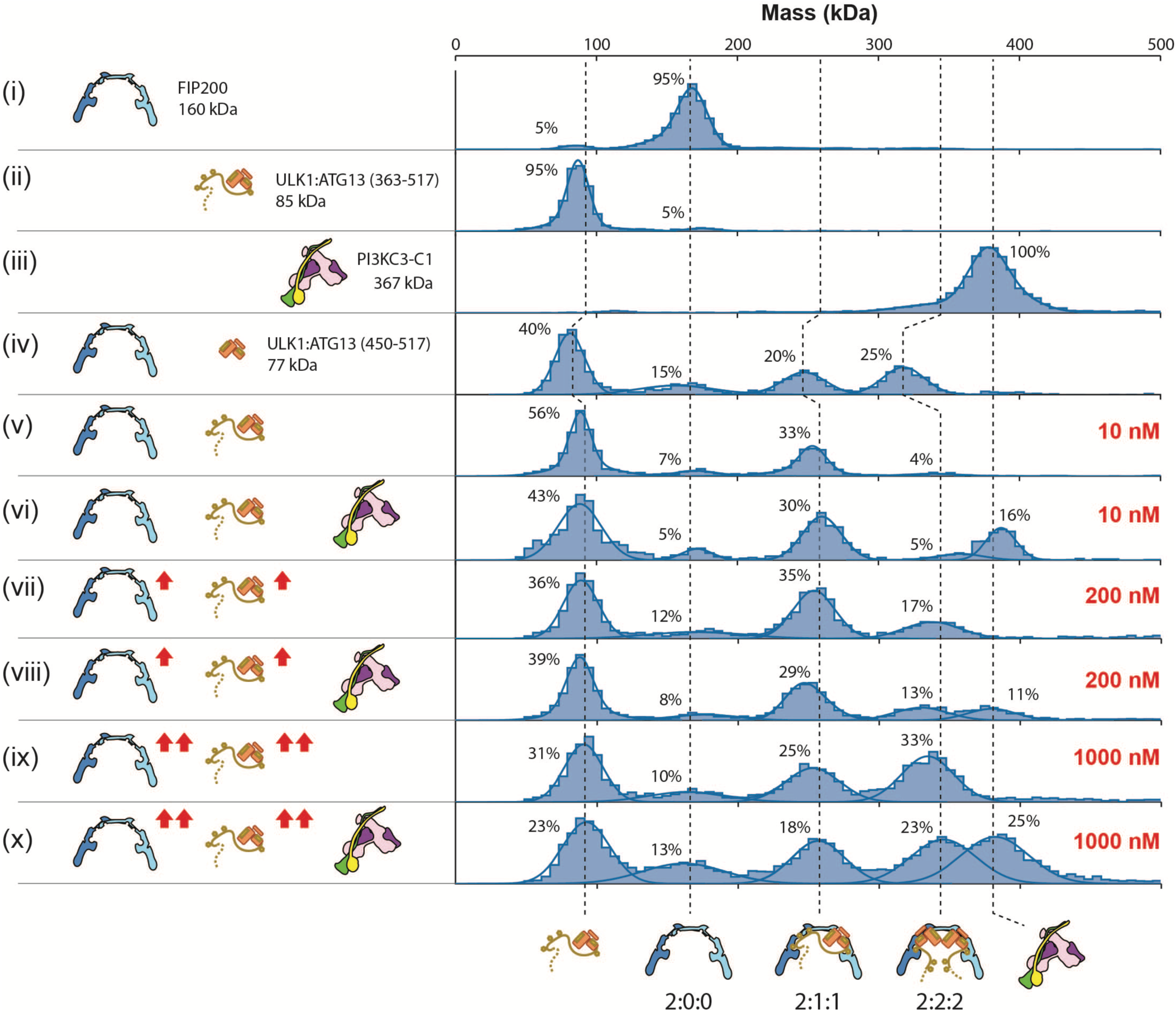
Concentration-dependent stoichiometry shift of ULK1C. Mass photometry measurements of (i) FIP200^NTD^, (ii) ULK1^MIT^:ATG13 (363-517), (iii) full length PI3KC1-C1; (iv) FIP200^NTD^ with truncated ULK1^MIT^:ATG13 (450-517). Note the molecular weight shifted because of the truncation, (v and vi) FIP200^NTD^ with ULK1^MIT^:ATG13 (363-517) crosslinked at 10 nM in the absence or presence of 5 nM PI3KC3-C1. (vii and viii) FIP200^NTD^ with ULK1^MIT^:ATG13 (363-517) crosslinked at 200 nM in the absence or presence of 100 nM PI3KC3-C1. (ix and x) FIP200^NTD^ with ULK1^MIT^:ATG13 (363-517) at crosslinked 1000 nM in the absence or presence of 500 nM PI3KC3-C1. Solid curves represent fits to multiple Gaussian to estimate the average mass and the percentage of each population (Table S2). The percentage is calculated from the area of the peaks in the range of 0-500 kDa. The schematic drawing illustrates the setting of each measurement (left) and the predicted assemblies (bottom). The theoretical molecular weights of each assembly are shown next to the diagrams (i-iv, left). The crosslinking concentration of FIP200 and ULK1:ATG13 used for each measurement are indicated with red labels (v-x, right).

We sought to determine whether PI3KC3-C1 also promoted the formation of the 2:2:2 complex in solution. By mass photometry, we observed that the ULK1C core alone at 10 nM concentration formed a 2:1:1 complex (Figure 6-v), consistent with the SEC-MALS results. The presence of PI3KC3-C1 did not alter the stoichiometry in solution (Figure 6-vi). We hypothesized that PI3KC3-C1 might facilitate the membrane recruitment and increased local concentration of ULK1C in autophagy initiation, which might manifest in a cryo-EM experiment as recruitment and concentration of ULK1 at the air-water interface on EM grids. We found that upon increasing the concentration from 200 nM to 1 μM, abundant 2:2:2 complex appeared in solution, roughly co-equal to the 2:1:1 complex (Figure 6-ix). Addition of PI3KC3-C1 did not lead to a further enhancement of the 2:2:2/2:1:1 ratio in solution (Figure 6-x). This observation shows that mass action alone can drive the 2:1:1 to 2:2:2 state conversion, even in the absence of PI3KC3-C1 and for constructs containing ATG13 site 2.

## Discussion

Despite that ULK1C and PI3KC3-C1 are at the heart of human autophagy initiation ^35^, there have been major gaps in understanding how autophagy is switched on by these two complexes. The available structures of ULK1 complex components have been either too fragmentary or at inadequate resolution to draw mechanistic conclusions about the integrated regulation of the entire complexes. Beyond the ability of ULK1 to phosphorylate all four subunits of PI3KC3-C1^21–24^, it has been unclear how the activities of the two complexes are coordinated. Here, we obtained a cryo-EM reconstruction of the ULK1C core that permitted amino acid-level interpretation. We then found that ULK1C and PI3KC3-C1 form a physical complex mediated by ULK1C FIP200 and PI3KC3-C1 VPS15, BECN1, and ATG14 subunits. The ULK1C core binds full-length PI3KC3-C1 stably enough to yield a cryo-EM reconstruction. Our findings differ from a report indicating that ULK1C binds to a BECN1:ATG14 subcomplex of PI3KC3-C1 with full-length HORMA domain-containing ATG13 and ATG101 ^36^, which can be attributed to the use of the intact PI3KC3-C1 in this study versus the BECN1:ATG14 subcomplex used in ref. 36. In yeast, the Atg1 complex (the yeast counterpart of human ULK1C) and PI3KC3-C1 are also reported to be in direct physical contact, which mediates the recruitment of PI3KC3-C1 to the phagophore initiation site in yeast ^37^. The determinants of binding, however, differ between yeast and humans. Among other differences, Atg38 is important for PI3KC3-C1 recruitment in yeast ^37^, while we found that the human supercomplex assembles independently of the human Atg38 ortholog NRBF2. The same surface region of VPS15^HSD^ binds to both FIP200 and NRBF2 (Figure S7C) ^17^, therefore, the relationship between these interactions remains to be investigated further. The most important conclusions are that ULK1C and PI3KC3-C1 form a physical complex through interactions that involve the autophagy-specific ATG14 subunit of PI3KC3-C1 responsible for autophagy-specific functions.

We previously found that TBK1 contains a scaffold domain that closely resembles the arm of FIP200 involved in binding to VPS15 (Figure S2B), and TBK1 has been shown to bind directly to PI3KC3-C1 ^27^. Most, but not all, autophagy initiation pathways require ULK1/2. In OPTN-mediated Parkin-dependent mitophagy, TBK1 replaces the requirement for ULK1/2 ^27^. It will be interesting to determine if TBK1 uses its FIP200 arm-like domain to bypass the need for ULK1/2 in OPTN mitophagy. The role of the VPS15 pseudokinase domain in PI3KC3 function has been a mystery since its initial identification in yeast ^38^. We recently elucidated how VPS15 regulates the lipid kinase activity of PI3KC3 ^30^. Here, we established a role for VPS15 as the major bridge to FIP200 in ULK1C recruitment. The N-terminal predicted zinc-finger domain of ATG14 is crucial for autophagy initiation and the recruitment of PI3KC3-C1 to initiation sites ^5,15^. Yet the molecular function of the ATG14 N-terminal region in the context of PI3KC3-C1 has been elusive and it has not been visualized in previous PI3KC3-C1 structures. Here, we found that this region co-folds with the BECN1 N-terminal domain, and both wrap around the FIP200 proximal arm, consistent with and potentially explaining the function of this region in autophagy initiation. A role for BECN1^BH3^ in regulating autophagy via interactions with Bcl-1 is also well-established ^39–41^. Our observation that BECN1^BH3^ participates in ULK1 supercomplex formation suggests that BECN1^BH3^ could have a more direct function in autophagy initiation than previously appreciated.

With respect to the regulation of autophagy initiation, we observed a rearrangement of ULK1C from a 2:1:1 to a 2:2:2 complex. ULK1 kinase activity requires autophosphorylation at Thr180 ^42^, as does its yeast ortholog, Atg1 ^43^. Artificially induced dimerization of yeast Atg1 promotes its activation ^44,45^. The dimerization and subsequent autophosphorylation of receptor-linked tyrosine kinases (RTKs) is the central paradigm for kinase activation in growth factor signaling ^46^. These findings suggest that PI3KC3-C1- and ATG13-regulated ULK1 kinase dimerization could be an autophagic cognate of the dimerization-based RTK activation paradigm. Here, we observed that the presence of PI3KC3-C1 induced 2:2:2 conversion on cryo-EM grids, but not in solution. We established that mass action can drive 2:2:2 conversion in solution in the absence of PI3KC3-C1. We found that ATG13^IDR^ site 2 regulates 2:2:2 conversion both on cryo-EM grids and in solution, demonstrating a common molecular mechanism in both settings. In the supercomplex structure, the N-terminal portions of PI3KC3-C1 subunits BECN1 and ATG14 closely approach ATG13^IDR^ sites 1-2 (Figure S7D and S7E). ULK1 initiates autophagy in the context of puncta of > 30 molecules per cluster that are co-localized with the ATG14 subunit of PI3KC3-C1 ^47,48^.The elevated local concentration of ULK1 and associated molecules under conditions of co-recruitment and co-localization with PI3KC3-C1 would be expected to help drive formation of the 2:2:2 complex. Taking all of these observations together, we suggest that 2:2:2 conversion is a versatile mechanism for ULK1 activation that can integrate various driving inputs in distinct contexts (Figure 7).

**Figure 7.**
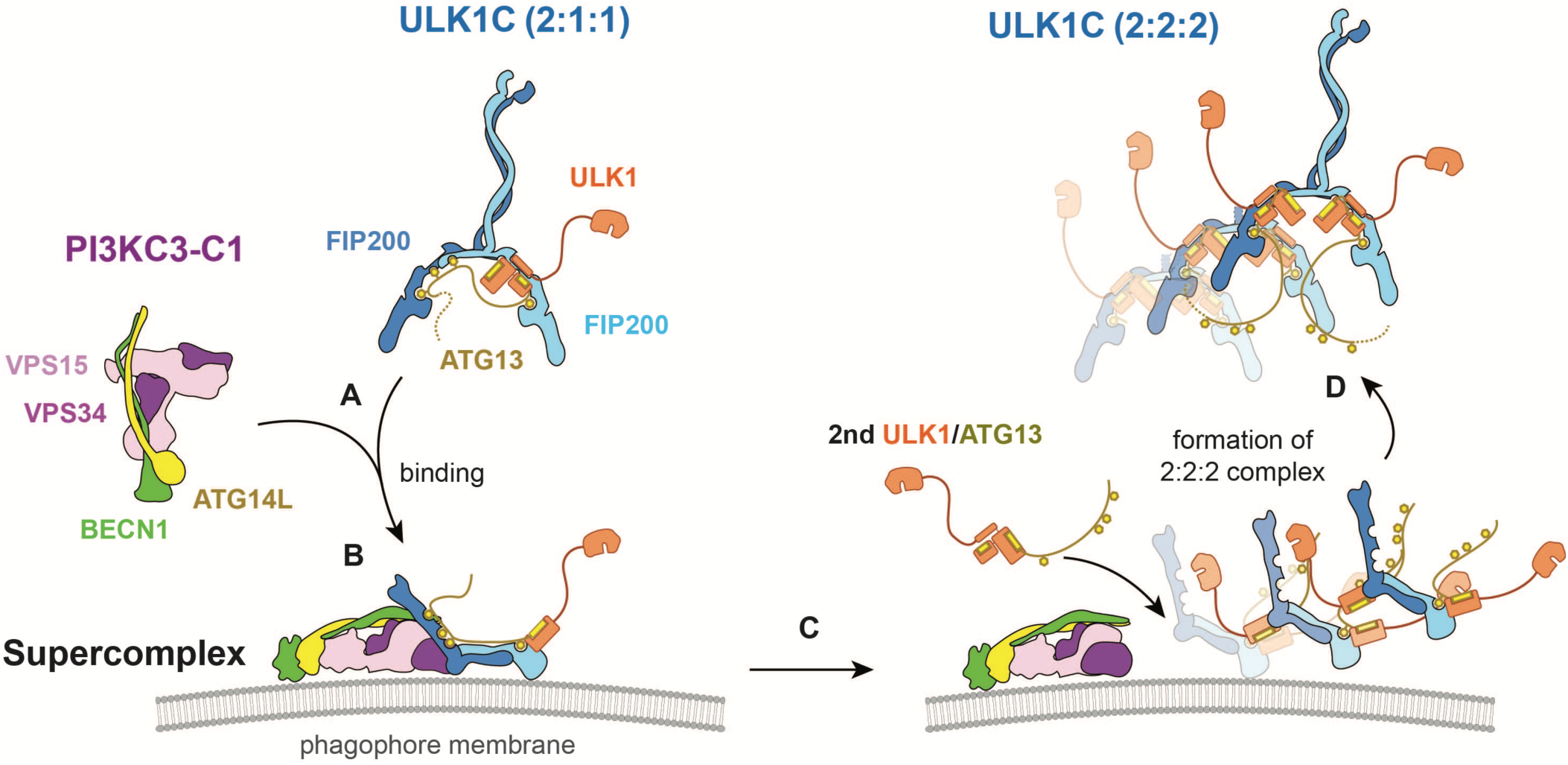
Model for ULK1 activation. (**A**) Binding and recruitment of the ULK1C to the phagophore membrane by PI3KC3-C1. (**B**) Formation of ULK1C:PI3KC3-C1 supercomplex. (**C**) Increase of the local concentration of ULK1C facilitated by PI3KC3-C1. (**D**) Formation of ULK1 (2:2:2) complex.

Given the large interface between PI3KC3-C1 and FIP200, the multiple contacts between the ATG13^IDR^ and FIP200, it seems likely that various upstream kinases and other signaling inputs could regulate formation of the ULK1C:PI3KC3-C1 supercomplex and the activated 2:2:2 ULK1C. While mTORC1 is one candidate to modulate these complexes, these mechanisms are in principle general and could be regulated by other kinases such as AMPK ^49^ or TBK1 ^50^, or by increased local concentration and colocalization driven by selective autophagy cargo receptor clustering ^51^ or condensate formation ^52^. The new activation-related interfaces identified in this study are also suggestive of new concepts for the therapeutic upregulation of autophagy. The concept of ULK1C:PI3KC3-C1 supercomplex formation at the heart of autophagy initiation thus puts both the signaling and therapeutic aspects of autophagy initiation into new perspective.

## Resource availability

### Lead contact

Further information and requests for resources and reagents should be directed to and will be fulfilled by the lead contact, James H. Hurley

### Materials availability

All unique regents generated in this study will be available from the lead contact upon request.

### Data availability

The cryo-EM maps were deposited in the Electron Microscopy Data Bank (EMDB) under accession codes EMD-40658 (ULK1C (2:1:1) core), EMD-45297 (ULK1C:PI3KC3-C1 supercomplex), EMD-40715 (ULK1C (2:2:2) core in the PI3KC3-C1 mixture), and EMD-40735 (ULK1C (2:2:2) core of the ATG13^450–517^ truncation mutant). The structural coordinates were deposited in the Protein Data Bank (PDB) under accession codes 8SOI (ULK1C (2:1:1) core), 9C82 (ULK1C:PI3KC3-C1 supercomplex), 8SQZ (ULK1C (2:2:2) core in the PI3KC3-C1 mixture), and 8SRM (ULK1C (2:2:2) core of the ATG13^450–517^ truncation mutant). Protocols were deposited in protocols.io. Plasmids developed for this study were deposited at Addgene.org. Raw data files for gel scans were uploaded to Zenodo.

## Acknowledgments

We thank A. Yokom and X. Shi for contributing to early stages of the project, and all members of the Hurley Lab, and D. Fracchiolla and others in Aligning Science Across Parkinson’s (ASAP) Team mito911 for advice and discussions. We thank D. Toso, P. Tobias and R. Thakkar for cryo-EM facility support, and E. Nogales for mass photometry. This research was funded by Aligning Science Across Parkinson’s [ASAP-000350] through the Michael J. Fox Foundation for Parkinson’s Research (MJFF) (to M.L. and J.H.H.), and National Institutes of Health (NIH) [R35GM136414] (to A.Y.).

## Author contributions

Conceptualization: J.H.H. Methodology: M.C., X.R., T.N.N., G.K., Y.Z. and A.C. Investigation: M.C., X.R., T.N.N., Y.Z. and A.C. Visualization: M.C. Supervision: M.L., A.Y. and J.H.H. Writing—original draft: M.C. and J.H.H. Writing—review and editing: All authors.

## Methods

### Key resource table

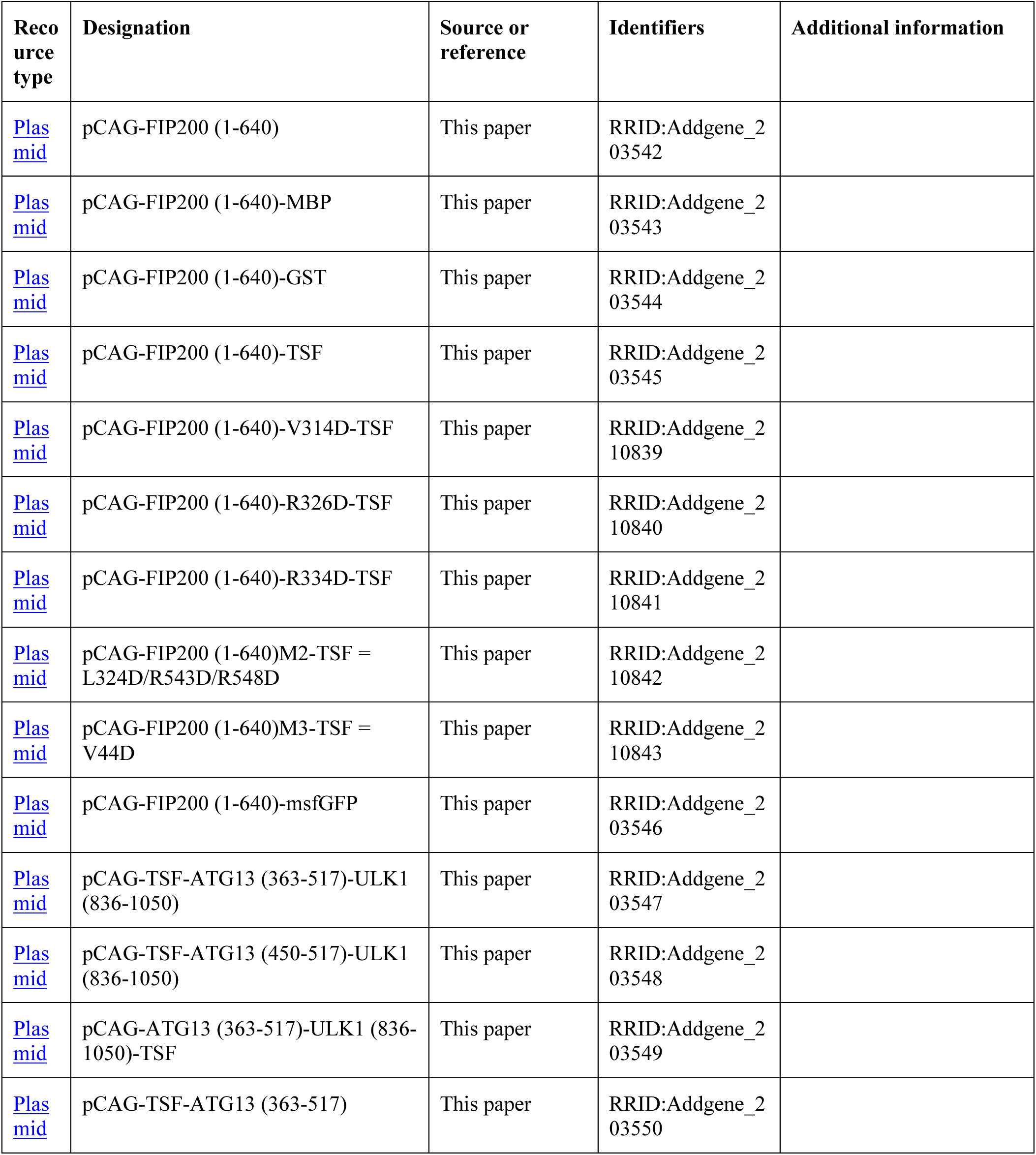
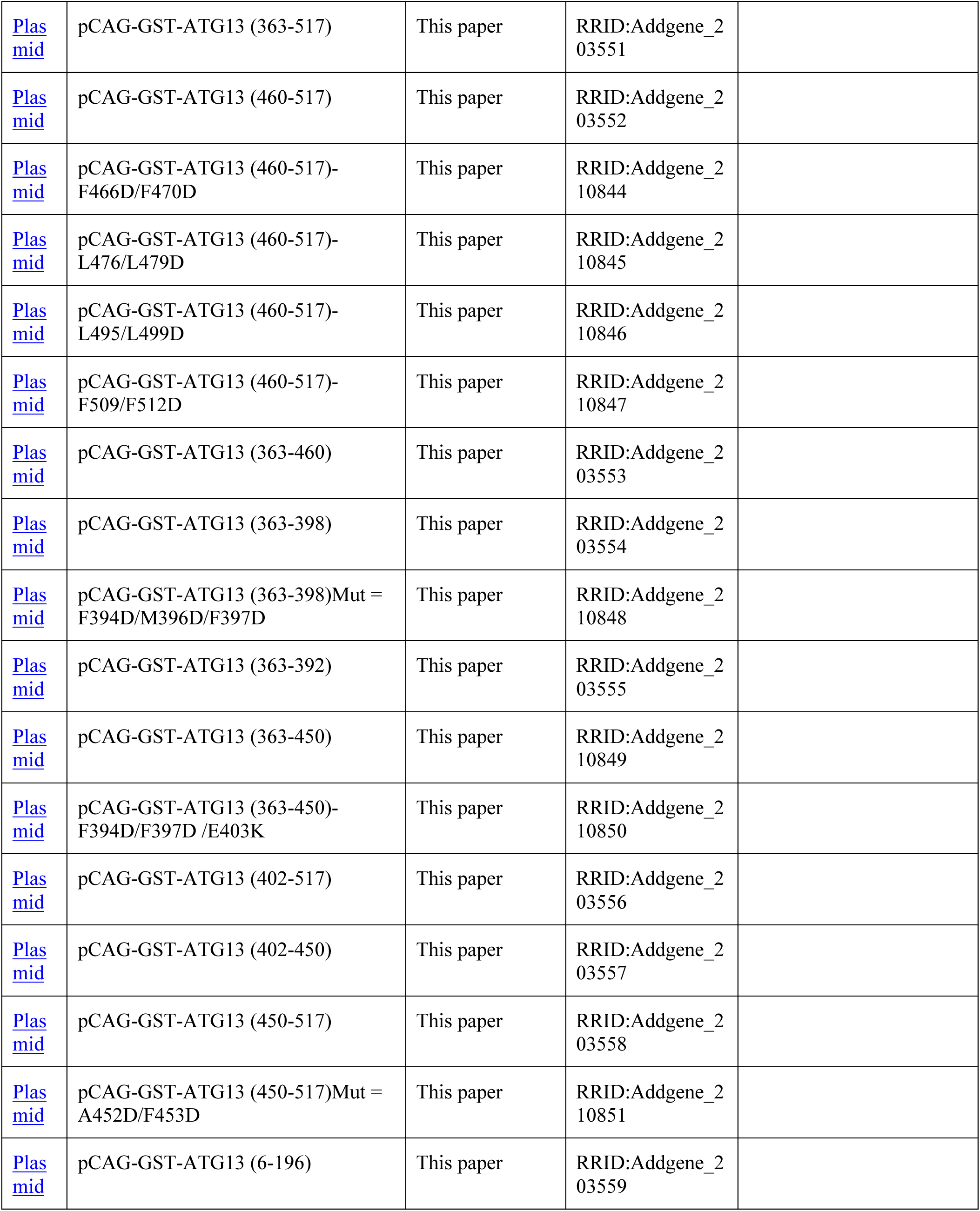
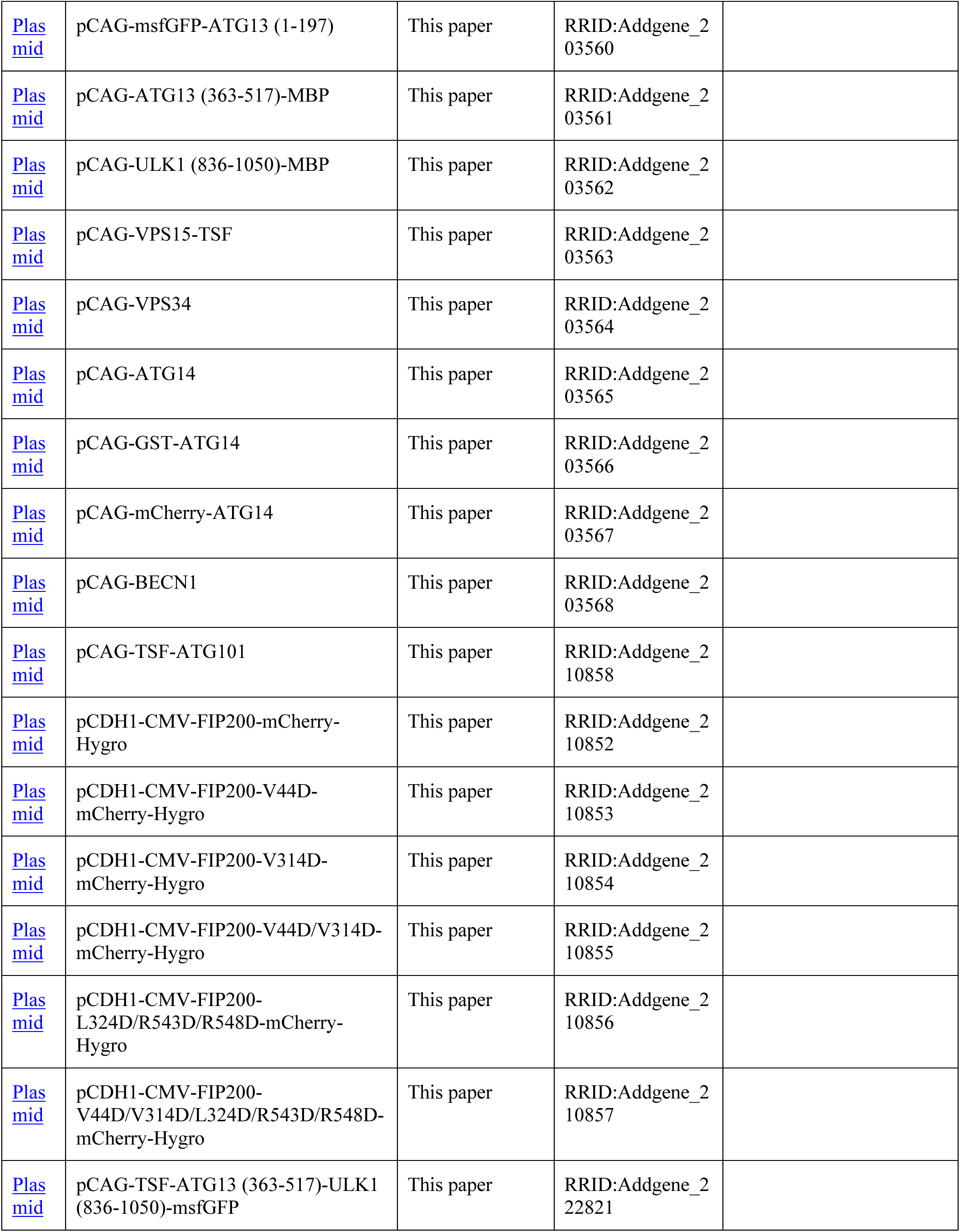
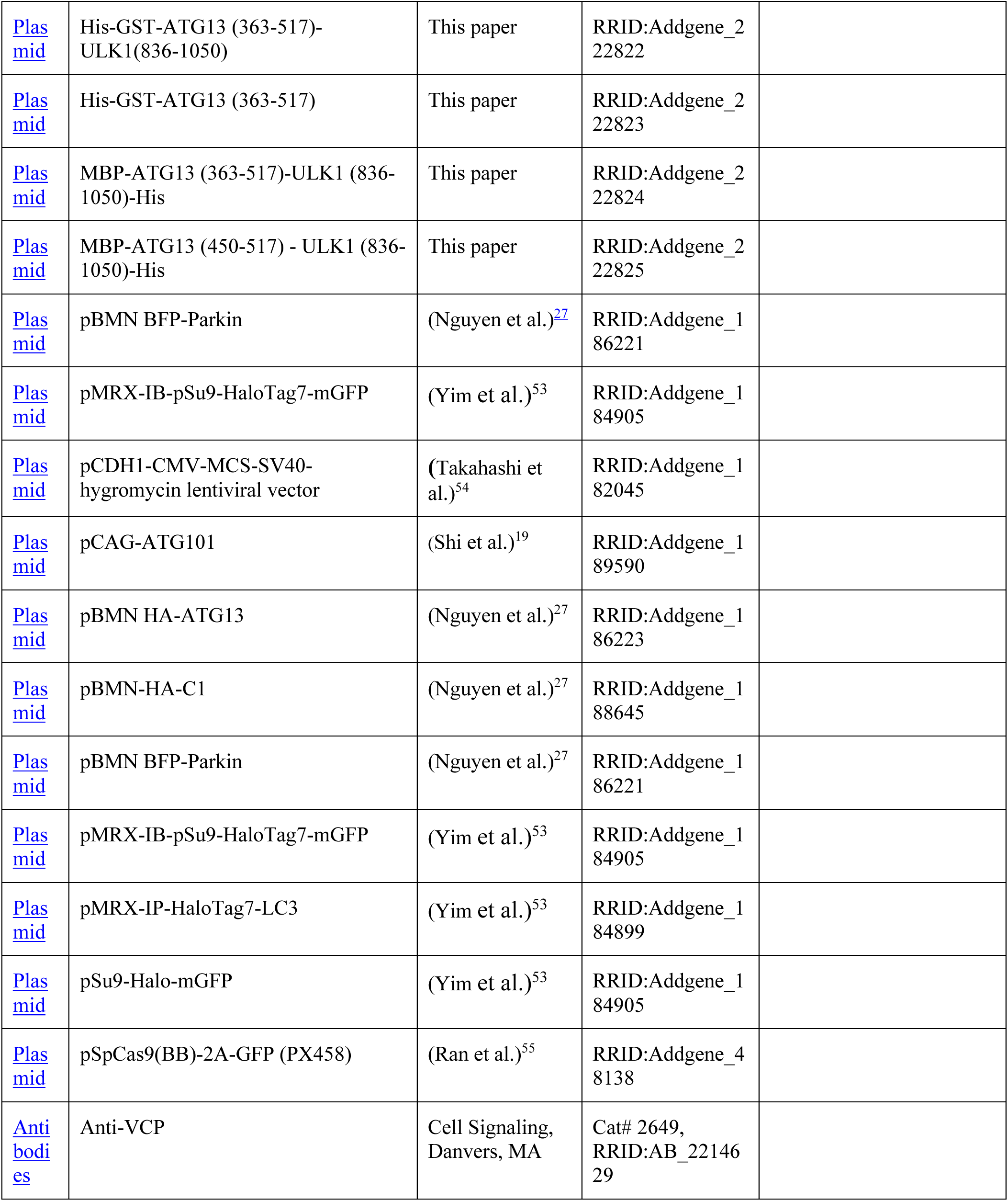
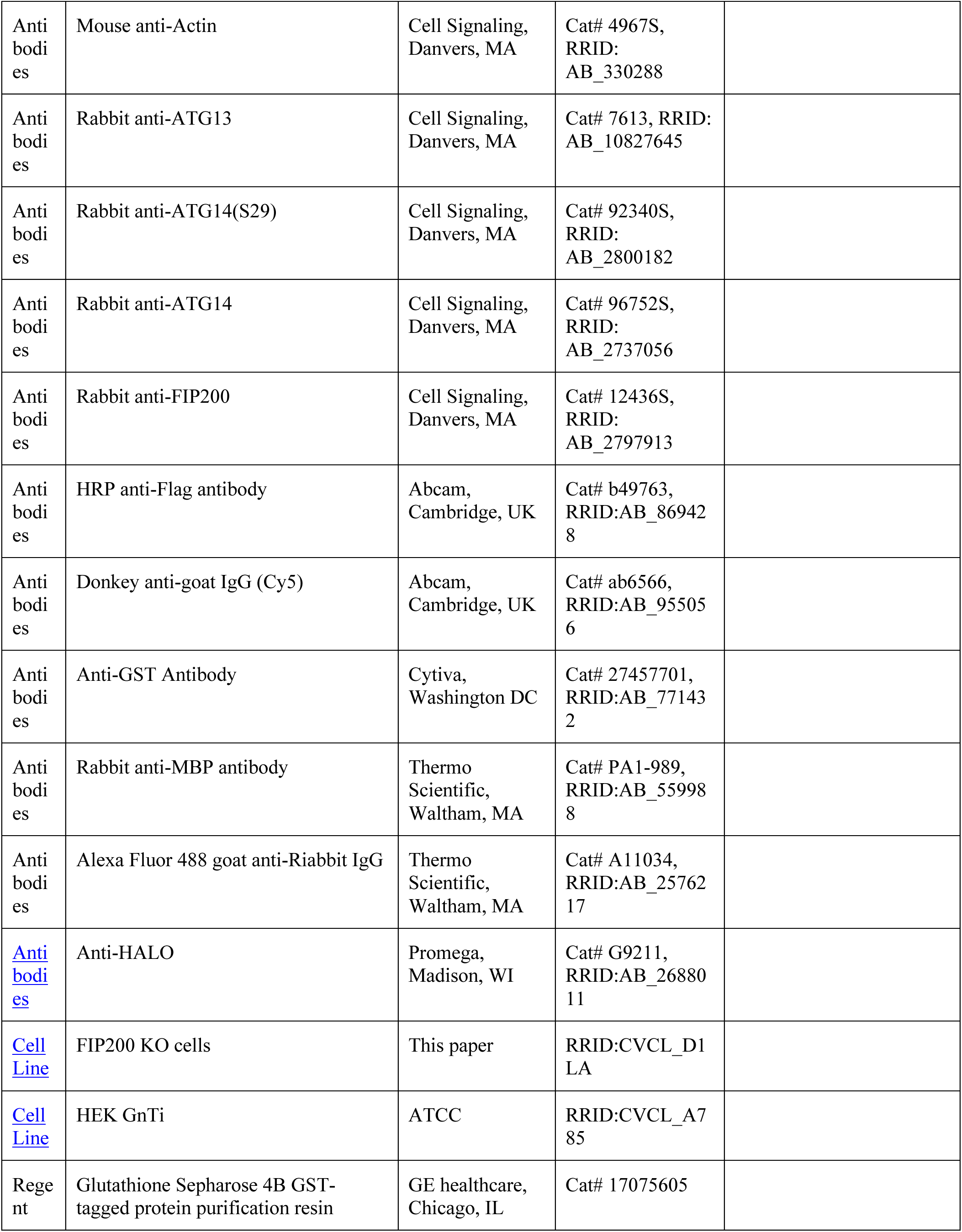
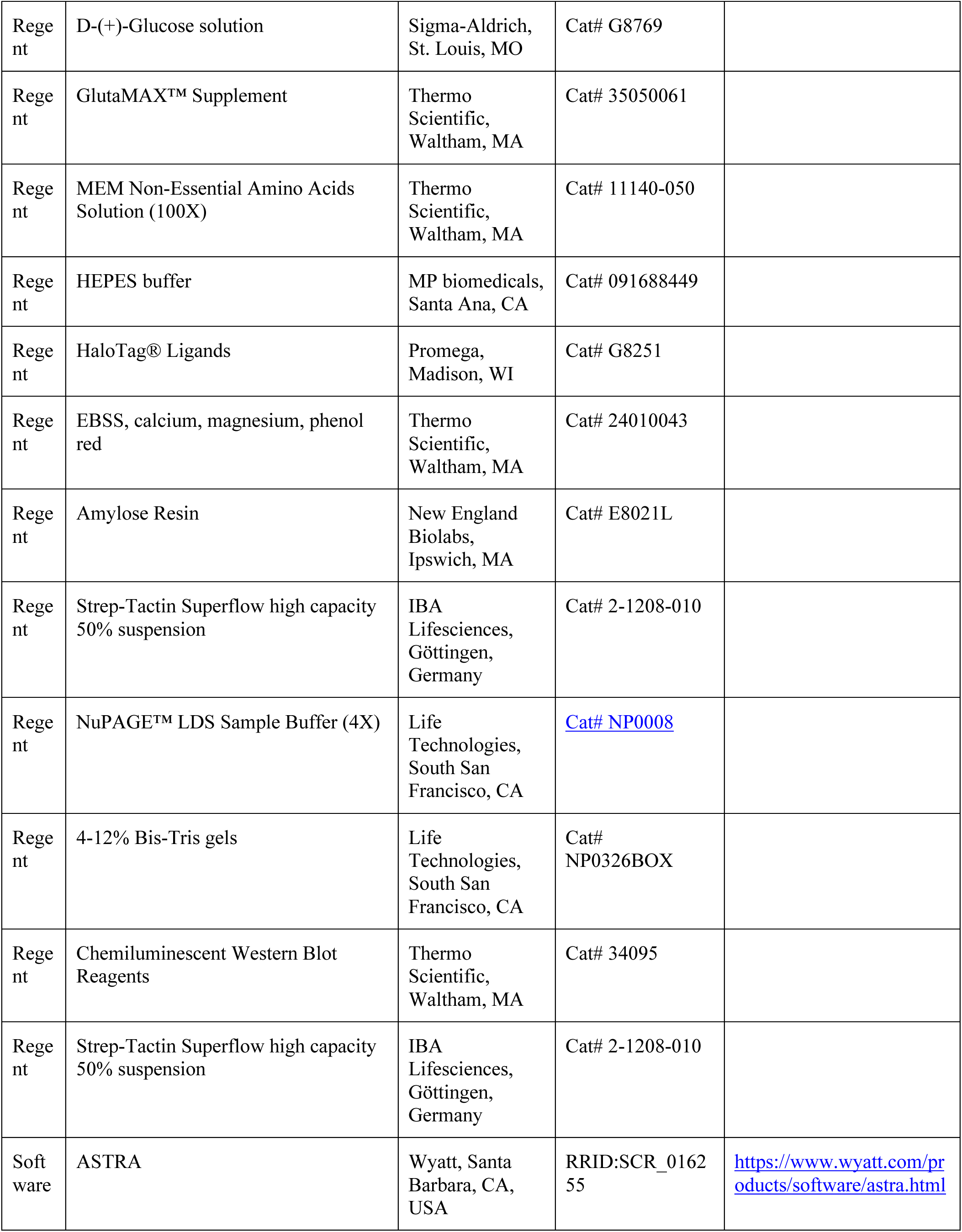
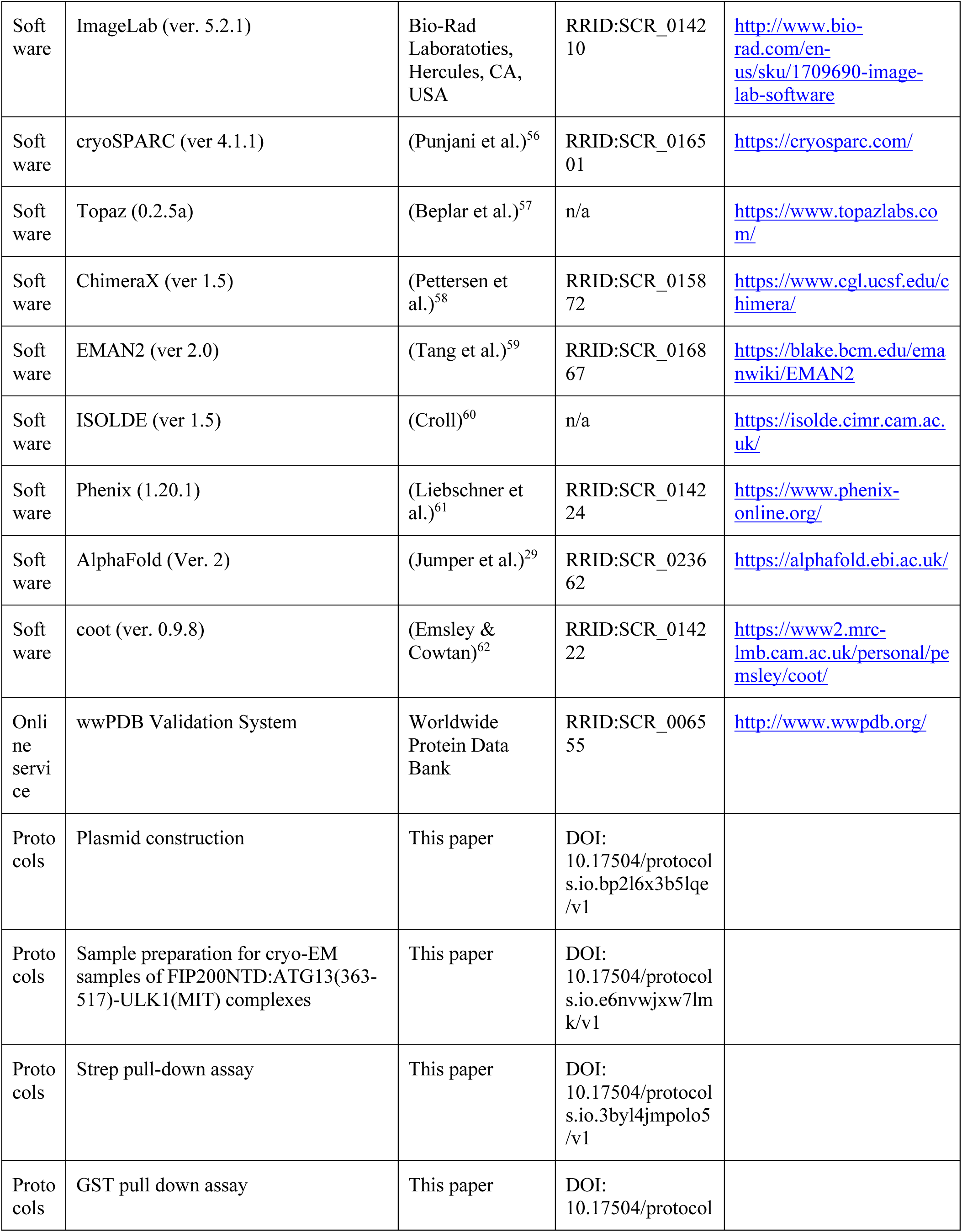
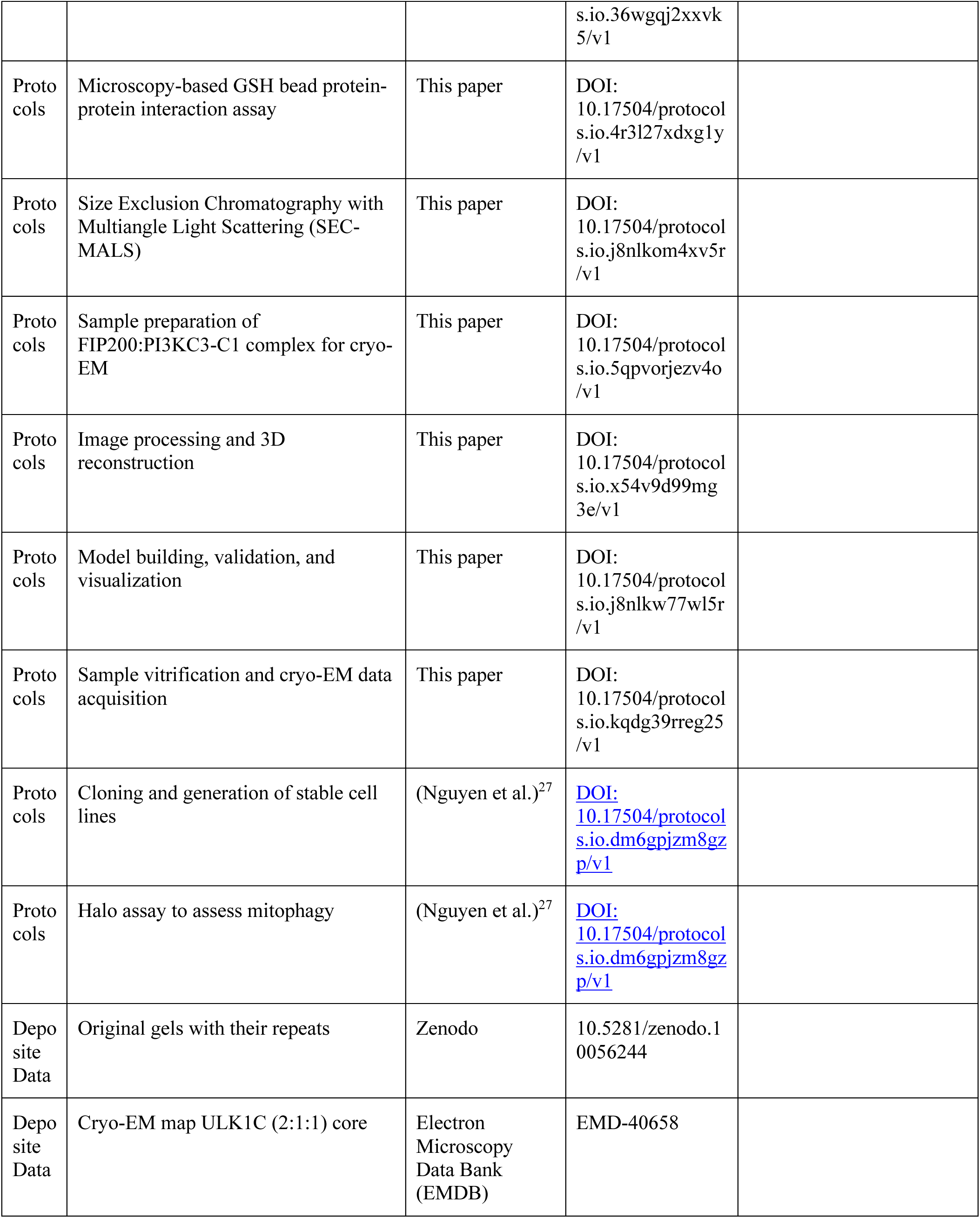
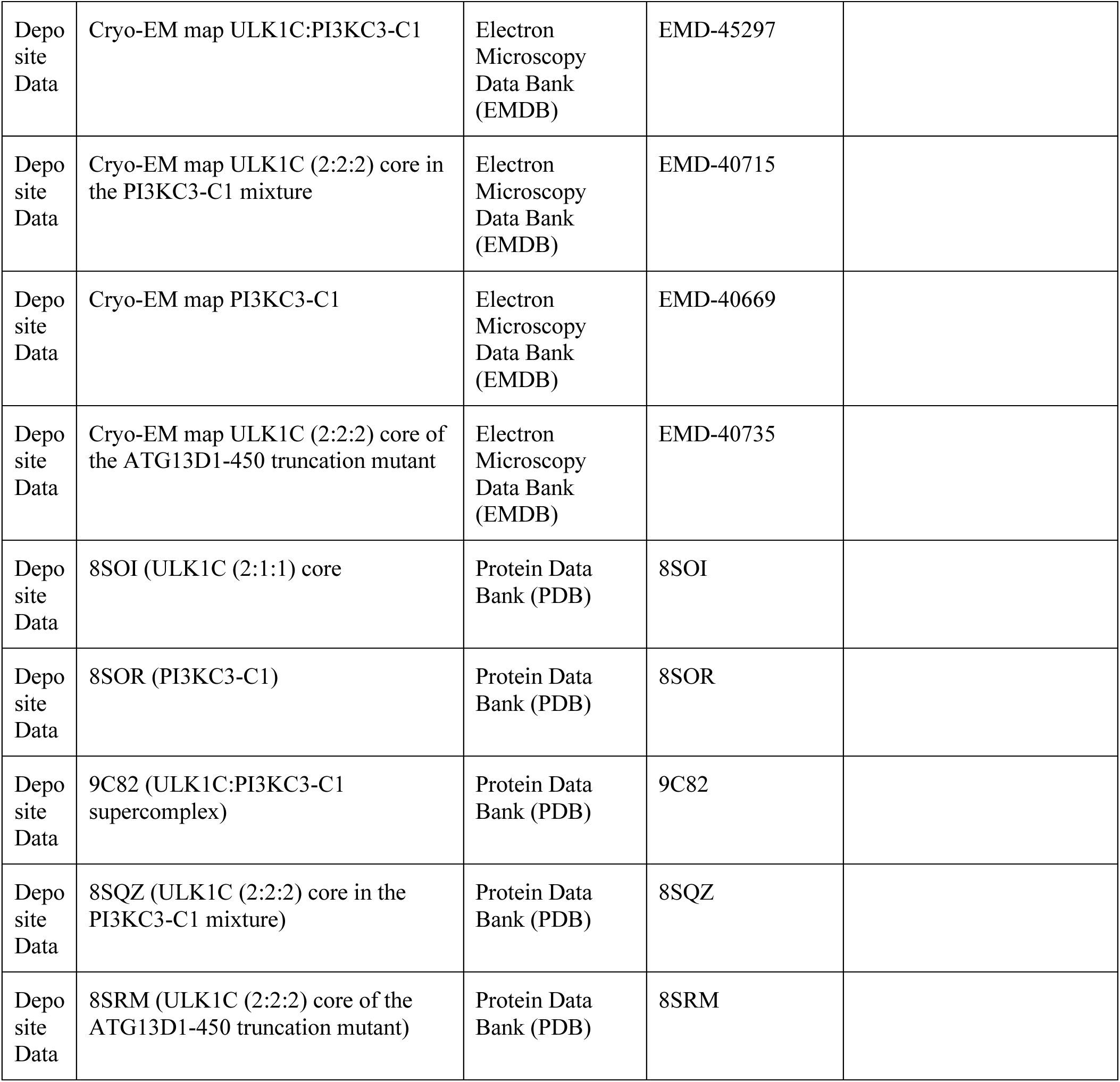

## Method details

### Plasmid construction

The sequences of all DNAs encoding components of human ULK1C were codon optimized, synthesized and then subcloned into the pCAG vector. The fusion construct of ATG13 C-terminal with ULK1^MIT^ domain was subcloned as a TwinStrep-Flag (TSF), TEV cleavage site, ATG13 (363-517) or (450-517), 5 amino acid linker (GSDEA) followed by ULK1 (836-1050) into the pCAG vector. Proteins were tagged with GST (Glutathione S-Transferase), MBP (Maltose-Binding Protein) or TSF for affinity purification, pull-down or GSH (Glutathione) beads assay. All constructs were verified by DNA sequencing. Details are shown in key resource table, STAR Methods.

### Protein expression and purification

For cryo-EM samples of FIP200^NTD^:ATG13^363–517^-ULK1^MIT^ complexes, pCAG-FIP200^NTD^ (1-640) was co-transfected with pCAG-TSF-ATG13^363–517^-ULK1^MIT^ (836-1050) using the polyethylenimine (PEI) (Polysciences) transfection system. FIP200^NTD^-MBP:ATG13^363–517^-ULK1^MIT^-TSF were co-expressed in HEK293 GnTi^-^ cells for the FIP200:PI3KC3-C1 cryo-EM study. Cells were transfected at a concentration of 2 × 10^6^/ml. After 48 hours, cells were pelleted at 500x g for 10 min, washed with PBS once, and then stored at −80°C. Cell pellets were lysed at room temperature for 20 min with lysis buffer (25 mM HEPES pH 7.5, 200 mM NaCl, 2 mM MgCl_2_, 1 mM TCEP, 10% Glycerol) with 5 mM EDTA, 1% Triton X-100 and protease inhibitor cocktail (Thermo Scientific) before being cleared at 17000x rpm for 35 min at 4 °C. The clarified supernatant was purified on Strep-Tactin Sepharose resin (IBA Lifesciences) and then eluted in the lysis buffer with 4mM desthiobiotin (Sigma). After His_6_-TEV cleavage at 4°C overnight, samples were concentrated then load onto a Superose 6 Increase 10/300 GL column (Cytiva) in the buffer of 25 mM HEPES pH 7.5, 150 mM NaCl, 1 mM MgCl_2_, 1 mM TCEP.

PI3KC3-C1 complex was expressed in HEK293 GnTi^-^ cells via PEI transfection from codon optimized pCAG-VPS15-TSF, pCAG-VPS34, pCAG-ATG14 and pCAG-Beclin-1. pCAG-GST-ATG14 or pCAG-mCherry-ATG14 was used in PI3KC3-C1 expression for GST pull-down or microscopy-based GSH bead assay. Cells were transfected at a concentration of 2 × 10^6^/ml and harvested after 48-hour post-transfection. Pellets were homogenized 20 times by Pyrex douncer (Corning) in lysis buffer with 25mM TCEP/proteinase inhibitors (Thermo Scientific), and then add 10% Triton X-100 stock to final 1% concentration. After rocking at 4 °C for 1 hour, lysates were clarified by centrifugation (17,000 x rpm for 40 min at 4 °C) and incubated with Strep-Tactin Sepharose (IBA Lifesciences) at 4 °C overnight.

After eluting with lysis buffer/4 mM desthiobiotin, samples were concentrated and then loaded onto a Superose 6 Increase 10/300 GL column (Cytiva) in 25 mM HEPES pH 7.5, 300 mM NaCl, 1 mM MgCl_2_, 25 mM TCEP. FIP200^NTD^ constructs used in Fig. 3, A and B were purified using strep-tactin Sepharose resin as described above, and then loaded onto a Superose 6 Increase 10/300 GL column (Cytiva). ATG101:ATG13N HORMA related proteins in Fig. 3, A and B were purified using strep resin and a Superdex 200 Increase 10/300 GL column (Cytiva) equilibrated in 25 mM HEPES pH 7.5, 150 mM NaCl, 1 mM MgCl_2_, 1 mM TCEP.

### Strep pull-down assay

FIP200^NTD^-TSF WT or mutants were co-transfected with GST-ATG13^363–517^ and/or ULK1^MIT^-MBP in 10 ml of HEK293 GnTi^-^ cells. The cells were harvested 48 hours after transfection. The pellets were homogenized in 0.5 ml of lysis buffer/protease inhibitors/1% TritonX-100, and clarified after 40,000g x 15 min. The lysate was incubated with 30 µl Strep-Tactin Sepharose resin (IBA-Lifesciences) at 4°C for 3 hours. The beads were washed four times, and then eluted in 50 µl lysis buffer/4 mM desthiobiotin. 18 ml eluent was mixed with lithium dodecylsulfate (LDS)/BME buffer, heated at 60°C for 5 min and subjected to SDS/PAGE gel. The gel was then stained with Coomassie brilliant blue G250. For loading input control, 0.3% lysate per sample was mixed with 1x LDS sample buffer (Life Technologies) containing 100 mM BME, boiled at 95 °C for 5min, and then analysed with 4-12% Bis-Tris gels (Life Technologies). Gels were electro-transferred to polyvinyl difluoride membranes (PVDF) and immunoblotted with various antibodies. FIP200-NTD-TSF lysate input was visualized using HRP anti-Flag antibody (Abcam, ab49763) and SuperSignal West Femto Maximum Sensitivity Substrate ECL kit (Thermo Scientific, 34095). GST-ATG13c lysate input was analysed using goat anti-GST antibody (Cytiva, 27457701) and Donkey anti-goat IgG (Cy5) (Abcam, ab6566). ULKc-MBP lysate was analysed using Rabbit anti-MBP antibody (Thermo Scientific, PA1-989) and Alexa Fluor 488 goat anti-Riabbit IgG (Invitrogen, A11034). The chemiluminescence or Fluorescence signals were detected by ChemiDocMP Imaging system (Biorad).

### Microscopy-based GSH bead protein-protein interaction assay

A mixture of 1 µM purified GST tagged protein and 500 nM purified fluorescent protein in total 70 µl volume was incubated with 9 µL pre-blocked Glutathione Sepharose beads (Cytiva) in a reaction buffer containing 25 mM HEPES at pH 7.5, 150 mM NaCl, 1 mM MgCl_2_ and 1 mM TCEP. After incubation at room temperature for 30 min, samples were mixed with additional 100 µL reaction buffer, and then transferred to the observation chamber for imaging. Images were acquired on a Nikon A1 confocal microscope with a Nikon Plan APO VC 20x/0.75 NA UV Microscope Objective. Three biological replicates were performed for each experimental condition.

### Size Exclusion Chromatography with Multiangle Light Scattering (SEC-MALS)

The purified FIP200^NTD^:ATG13-ULK1^MIT^ complexes were concentrated to 4-5 mg/ml. SEC-MALS experiments were performed using an Agilent 1200 HPLC system (Agilent Technologies, Santa Clara, CA), coupled to a Wyatt DAWN HELEOS-II MALS instrument and a Wyatt Optilab rEX differential refractometer (Wyatt, Santa Barbara, CA). For chromatographic separation, a WTC-050S5 size-exclusion column (Wyatt) with a 40 μl sample loop was used at a flow rate of 0.3 ml/min in the buffer of 25 mM HEPES pH 7.5, 200 mM NaCl, 1 mM MgCl_2_, 2 mM TCEP. The outputs were analyzed by the ASTRA V software (Wyatt). MALS signals, combined with the protein concentration determined by refractive index, were used to calculate the molecular mass of the complex.

### Sample preparation of FIP200:PI3KC3-C1 complex for cryo-EM

FIP200^NTD^-MBP:ATG13^363–517^-ULK1^MIT^-TSF (final 5 µM) was mixed with PI3KC3-C1-TSF complex at 1.5:1 molar ratio in total 200 µl volume, rocking at 4°C overnight. Next day, the sample was incubated with 50 µl Strep-Tactin Sepharose resin (IBA Lifesciences) at 4°C for 3 hr. After one wash, the beads were eluted in 50 µl of 10 mM d-Desthiobiotin with the buffer of 25 mM HEPES pH 7.5, 200 mM NaCl, 1 mM MgCl_2_, 10 mM TCEP.

### Sample vitrification and cryo-EM data acquisition

For cryo-EM sample preparation, 3 μl of protein solution was applied onto a grid freshly glow-discharged in PELCO easiGlow system (Ted Pella). In-house graphene grids were prepared from Trivial Transfer Graphene sheets (ACS Material) and QUANTIFOIL R2/1 mesh 300 gold (Electron Microscopy Sciences) by following a protocol introduced by Ahn et., al ^63^. The graphene grids were used for the ULK1C (2:1:1) core sample, and holey carbon grids (QUANTIFOIL R1.2/1.3 or R2/1 mesh 300, Electron Microscopy Sciences) were sufficient for the other samples. The samples were vitrified with a Vitrobot cryo-plunger (Thermo Fisher Scientific) in plunging conditions optimized previously. 0.05%(w/v) of n-Octyl-Beta-D-Glucopyranoside was added in the sample solution as a surfactant before vitrification.

The datasets of the ULK1C (2:1:1) core, the ULK1C:PI3KC3-C1 mixture and the ULK1C:PI3KC3-C1 pull-down were recorded at a 300 kV Titan Krios microscope (Thermo Fisher Scientific) equipped with X-FEG and energy filter set to a width of 20 eV. Automated data acquisition was achieved using SerialEM ^64^ on a K3 Summit direct detection camera (Gatan) at a magnification of 81,000x and a corresponding pixel size of 0.525 Å at the super-resolution mode and a defocus range of −0.8 to −2.0 μm. The ULK1C:PI3KC3-C1 pull-down sample was collected at 0°, 20°, and 30° tilted specimen stage.

Image stacks with 50 frames was collected with a total dose of 50 e/Å^2^. The dataset of the ULK1C (2:2:2) core was recorded at a 200 kV Talos Arctica microscope (Thermo Fisher Scientific) equipped with the K3 Summit camera in a super-resolution correlated-double sampling mode. The magnification and the pixel size were 36,000x and 0.5575 Å at the super-resolution mode, respectively. Other details of the dataset collection are summarized in Table S1.

### Image processing and 3D reconstruction

The datasets were processed by following the workflow in cryoSPARC ^56^. In brief, the super-resolution video stacks were motion-corrected and binned 2x by Fourier cropping using Patch Motion Correction. Contrast transfer function determination was done by Patch CTF Estimation, followed by manual removal of the outlier micrographs base on the estimated defocus and resolution value. Single particles were automatically picked by Topaz ^57^ based on a manually trained model and extracted with a window size that is 1.5 times larger than the target particle and further binned to 2-4x to facilitate the following processing. Two-dimensional (2D) classification was then used for removing obvious junk particles for the following classification. The initial models were obtained by using *ab initio* reconstruction. In case the reconstruction job failed to give a healthy initial model, the classes displaying high-resolution features in the 2D classification step were selected and used for *ab initio* reconstruction. Further classification was done at the 3D level by multiple rounds of heterogeneous refinement until a clean substack was obtained. The particles were re-extracted with the refined coordinates on micrographs at the original bin 2x pixel size and used for homogeneous refinement for multiple rounds until the final resolution converges. To further improve the quality of the map, local refinement was applied to the datasets of the ULK1C (2:1:1) core and the ULK1C:PI3KC3-C1 pull-down sample. Masking areas were decided by 3D Variability or 3D Flex and the masks were created by UCSF ChimeraX ^58^ and Volume Tools. Each local map was aligned to the consensus map and composed by using EMAN2 ^59^. The composed maps were then used for model building. The details of data processing are summarized in Table S1.

### Model building, validation, and visualization

The *in silico* models of the ULK1C (2:1:1) core and the full length PI3KC3-C1 were generated by AlphaFold2 prediction ^29^. The resolution of the observed maps enabled amino acid sequence assignment. The primary and secondary structure of the predicted models agrees well with the EM maps. A flexible model fitting by using the real-time molecular dynamics simulation-based program ISOLDE ^60^ implemented in the visualization software UCSF ChimeraX was performed, followed by iteratively refinement by using the model editing software Coot ^62^ manually and real-space refinement in Phenix ^61^ automatically.

The two ULK1C (2:2:2) core models were created by aligning two copies of FIP200^NTD^: ATG13^363– 517^:ULK1^MIT^ sub-trimeric complex to the FIP200 dimerization domain, followed by flexible model fitting with using ISOLDE in ChimeraX. The side chains were removed because of the moderate resolution of these maps. The ULK1C:PI3KC3-C1 supercomplex coordinates were generated by fitting of the individual structures of the ULK1C (2:1:1) and the PI3KC3-C1 to the cryo-EM map. The side chains were removed. The quality of the models was validated by using the validation tools in Phenix and the online validation service provided by wwPDB ^65,66^. The details of the model quality assessment are summarized in Table S1. All the figures and videos were made using UCSF ChimeraX.

### AlphaFold2 model prediction

Models of the FIP200^NTD^:ATG13^MIT^:ULK^MIM^ core domain were generated using the ColabFold implementation of AlphaFold2 ^29,67–70^. The default MMseqs2 ^71^ pipeline was employed to generate and pair MSAs for each protein complex predicted. We predicted the following complexes: dimeric FIP200(1-640), ULK1(828-1050), and ATG13(363-517); dimeric FIP200(1-640), ATG13(363-450); dimeric FIP200(1-640), ATG13(450-517), and ULK(828-1050); monomeric FIP200(1-640), and ATG13(363-450); and, monomeric FIP200(1-640), ATG13(450-517), and ULK(828-1050). In each case, the models were assessed based on the global predicted template modeling score (pTM), and the interfacial predicted template modeling (iPTM) score, with iPTM > 0.5 meriting further inspection. The predicted contacts between ATG13, FIP200, and ULK1 were manually inspected in UCSF ChimeraX, and the local predicted local distance difference test (pLDDT) score of the contacting residues was used in conjunction with an assessment of the physiological environment of the interfaces to judge the quality of the predicted residue contacts. The highest-scoring local pLDDT interfaces were used for figure-making and interpretation of hydrogen deuterium exchange data.

### Generation of knockout lines using CRISPR/Cas9

CRISPR guide RNAs (gRNAs) targeting FIP200 (5’-TATGTATTTCTGGTTAACACTGG-3’ and ATG13 (5’-CCGCGAGTTTGATGCCTTTG-3’ ^72^ and 5’-TTGCTTCATGTGTAACCTCTGGG-3’) were cloned into BbsI-linearised pSpCas9(BB)-2A-GFP vector ^55^ (a gift from Feng Zhang; Addgene plasmid # 48138) ^55^ using Gibson Cloning kit (New England Biolabs). gRNA containing constructs were then sequence-verified and transfected into WT HeLa cells with X-tremeGENE 9 (Roche) overnight and GFP-positive single cells were sorted by fluorescence activated cell sorting (FACS) into 96 well plates. Single cell colonies were left to grow and screened by immunoblotting for the loss of the targeted gene product.

### Cloning and generation of stable cell lines

pBMN-HA-WT ATG13 was previously described (Addgene #186223) ^73^. cDNAs encoding for ATG13 mutants (site2=F394D/M396D/F397D/E403K; site 4 = F449D/A452D/F453D/S454A and site 5 = L495D/L499D; site 2/4/5) were synthesized from Integrated DNA Technologies (IDT) and cloned into pBMN-HA-C1 (Addgene #188645) using Gibson Cloning kit (New England Biolabs). Generation of stable cell lines were previously described ^53^ and available at protocols.io (doi.org/10.17504/protocols.io.dm6gpjzm8gzp/v1). pMRX-IP-HaloTag7-LC3 (Addgene #184899) and HA-tagged ATG13 variants were stably expressed in *ATG13* KO cells using retroviral transduction.

### Halo assay to quantify starvation induced autophagy

We measured autophagic flux using the previously described method ^53,74^. In brief, 350k-400k cells were seeded the day before treatment in 6 well plates. Cells were treated with 50 nM TMR ligand (Promega) in growth media [DMEM with 10% FBS, 4.5 g/l Glucose (Sigma), 1x GlutaMAXTM (ThermoFisher), 1x NEAA (ThermoFisher), 25 mM HEPES. After 15 min, cells were washed 3 times with 1 x DPBS and treated with EBSS (Gibco) for indicated time periods. Following treatment, samples were analysed via immunoblotting as described below.

### Immunoblotting

Following feeding and treatment periods, the cells were washed with ice-cold 1x PBS, harvested using cell scrapers and lysed in lysis buffer (1x LDS sample buffer (Life Technologies) containing 100 mM dithiothreitol (DTT; Sigma)). Samples were boiled at 99 °C with shaking for 7 min and 25 μg (for Halo assays) and 70 μg (for the rest of the blots) of protein per sample was analysed with 4-12% Bis-Tris gels (Life Technologies) according to manufacturer’s instructions. Gels were electro-transferred to polyvinyl difluoride membranes (PVDF) and immunoblotted with antibodies against indicated antibodies. For quantification, band intensities were measured using ImageLab 5.2.1 (BioRad). Statistical significance was calculated from at least three independent experiments using two-way ANOVA. Error bars are means ± SD (standard deviation). The procedure for immunoblotting is described in detail in https://dx.doi.org/10.17504/protocols.io.dm6gpjzm8gzp/v1

### Mass photometry

High-precision coverslips (Azer Scientific) were cleaned by alternating between isopropanol and water three times in a bath sonicator, each 3 min, followed by air drying. The gasket was similarly cleaned with isopropanol and water three times, without sonication, then air-dried. A total of 19 µL of mass photometry buffer (30 mM HEPES pH 7.4, 5 mM MgSO4 and 1 mM EGTA) was added to the well for autofocus. The protein sample was then diluted in the mass photometry buffer in the wells to a concentration of 5–20 nM before data collection. If crosslinking was performed, 1 μL of 0.1% (v/v) glutaraldehyde in mass photometry buffer was added to 9uL protein sample and incubated for 10 minutes. The reaction was then quenched by adding 10 μL of 1M Tris-HCl pH 7.4. All the samples were diluted to 10 nM for the measurement of mass photometry. Protein contrast counts were acquired using a Refeyn TwoMP mass photometer, with three technical replicates. Mass calibration was performed using a standard mix of conalbumin, aldolase, and thyroglobulin. Mass photometry profiles were analyzed and fitted to multiple Gaussian peaks, with the mean, standard deviation, and percentages calculated using DiscoverMP software (Refeyn).

## Supplemental information

Tables S1 and S2 Figures S1-S8

---

**Figure S1.**
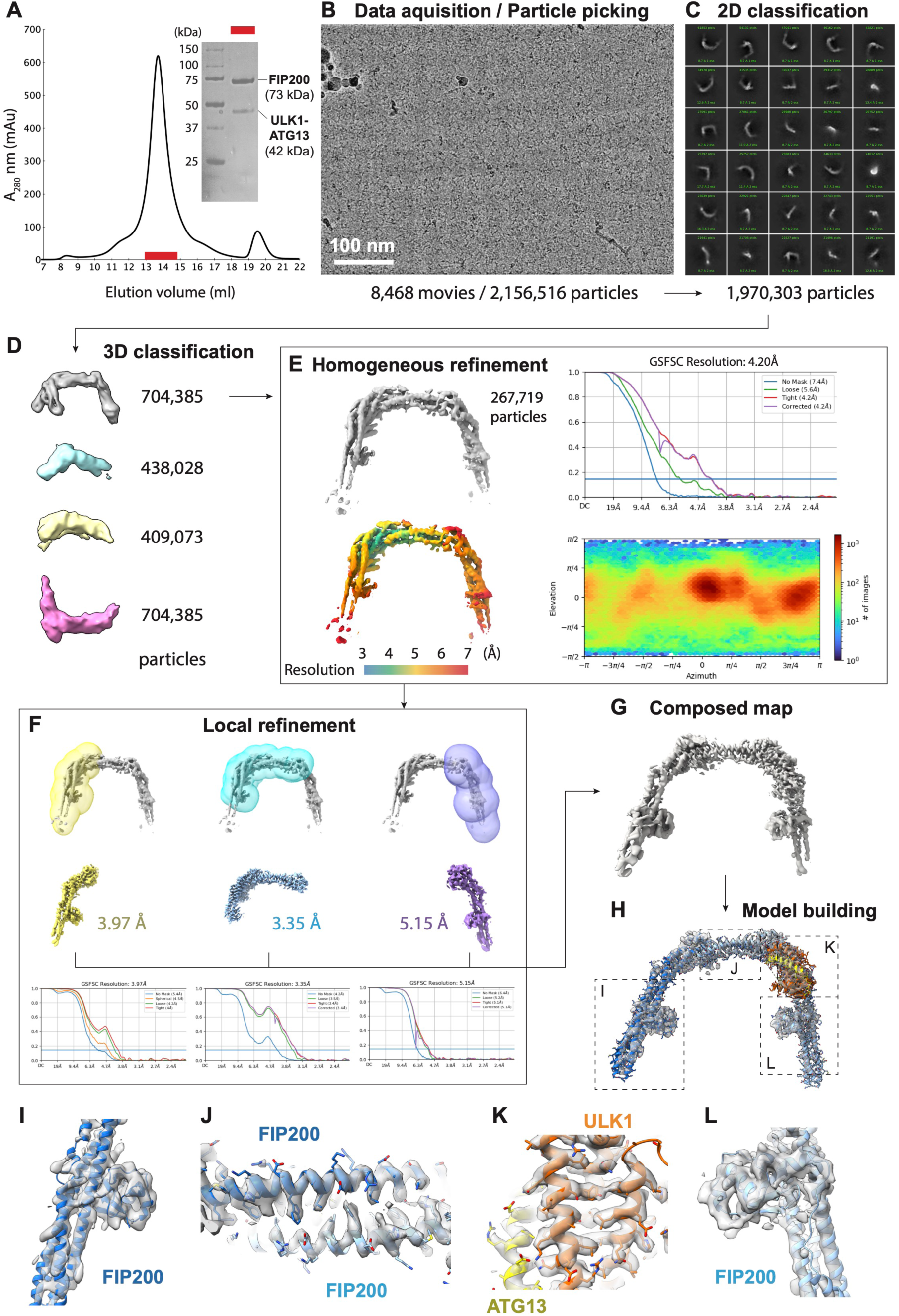
Cryo-EM sample preparation, image acquisition, data processing, and model building of ULK1C (2:1:1) core, related to Figure 1. (**A**) Size exclusion chromatography (SEC) profile of the ULK1C core. The inset shows an SDS-PAGE of the peak (red bar). (**B**) A representative cryo-EM micrograph of the ULK1C core. (**C**) Representative 2D class averages. (**D**) Result of the first round of 3D classification. (**E**) Global refinement of the final substack cleaned by multi-rounds of 3D classification. The FSC, local resolution, and angular distribution are shown accordingly. (**F**) Masks for local refinement and the corresponding maps and FSCs. (**G**) Final composed map for model building. (**H**) Overview of the model building. The map is contoured at 12σ. (**I to L**) Close-up views of the map superposed with the models. Each position is indicated in (**H**).

**Figure S2.**
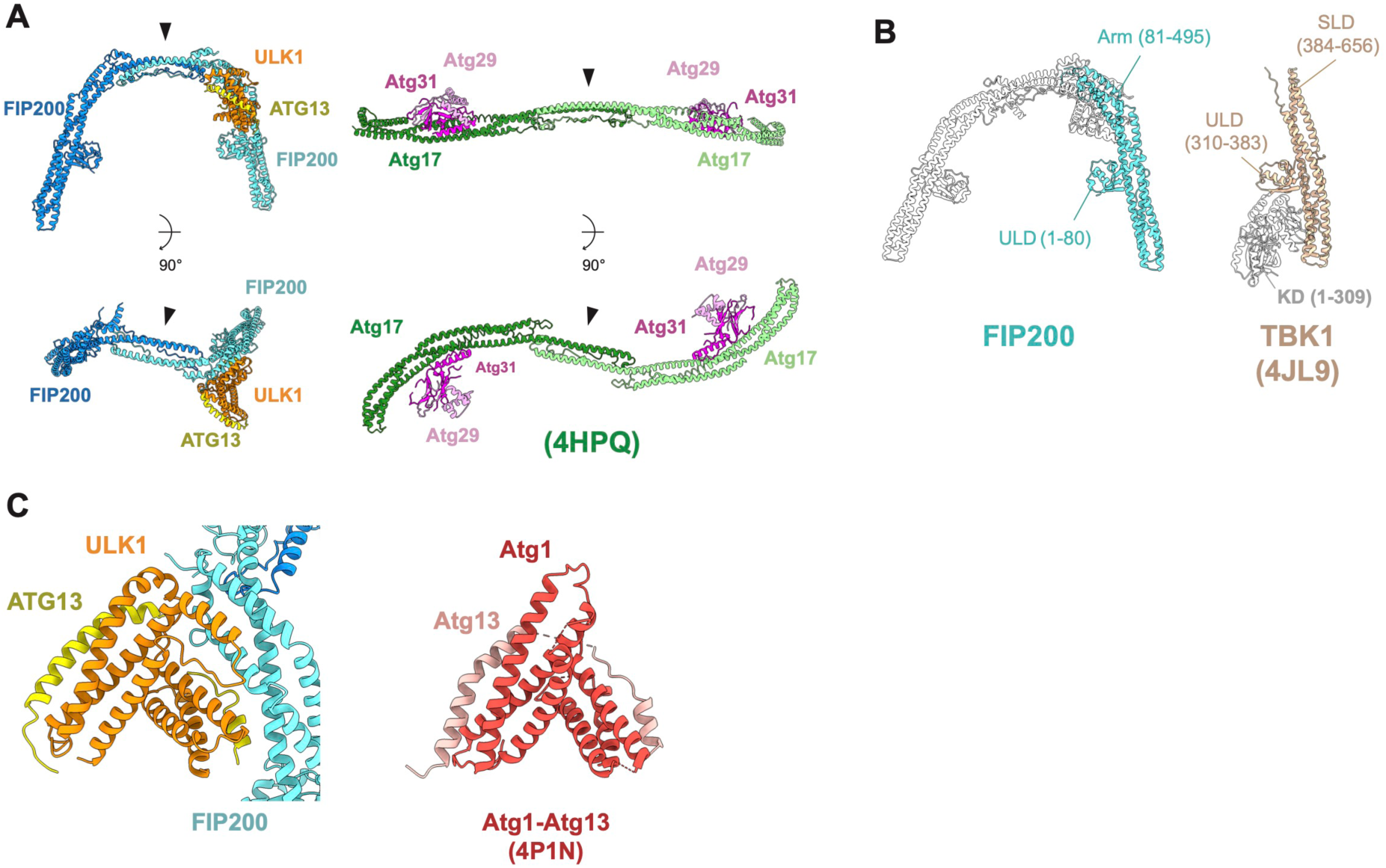
Structural comparison of the ULK1C (2:1:1) core with related proteins, related to Figure 1. (**A**) Structural comparison of the FIP200 with its yeast homolog Atg17 (PDBID: 4HPQ). The dimerization domains are indicated with arrows. (**B**) Structural comparison of the FIP200 with TBK1 (PDBID: 4JL9). (**C)** Structural comparison of the ULK1-ATG13 heterodimer with its yeast homolog Atg1-Atg13 (PDBID: 4P1N).

**Figure S3.**
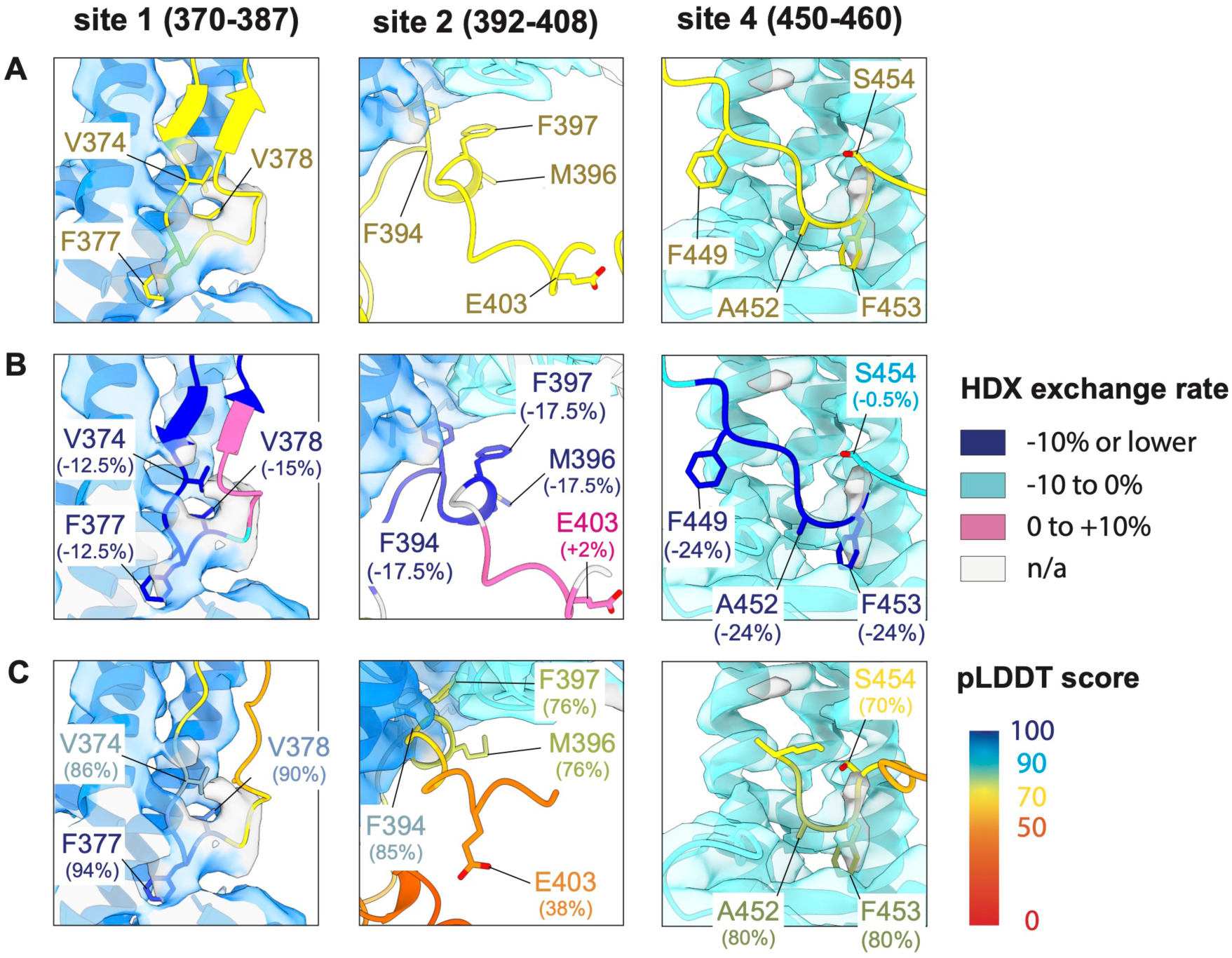
Assignment of the ATG13 binding sites, related to Figure 2. (**A**) Close up views of the binding site 1, 2, and 4 of the ULK1C core (2:1:1). The map is contoured at 11σ. The AlphaFold2 prediction model was created using dimeric FIP200(1-640), ULK1(828-1050), and ATG13(363-517), and superposed on the cryo-EM structure by aligning the FIP200 molecule. ATG13 is shown in yellow, while the other molecule of the predicted model are not shown. The key binding residues are indicated. (**B**) Same close up views of the binding site 1, 2, and 4. The ATG13 molecule is colored based on the HDX exchange rate (−10% or lower: blue; −10% to 0%: cyan; 0% to +10%: pink; n/a: gray) reported previously (*50*). The HDX exchange rate for each key residue is shown. (**C**) Same close up views of the binding site 1, 2, and 4. For optimizing the predicted local distance difference test (pLDDT) score, AlphaFold2 prediction models were created using monomeric FIP200(1-640) with ATG13(363-450) (left and middle), and monomeric FIP200(1-640) with ATG13(450-517) (right). The ATG13 molecule is colored based on the pLDDT score. (100%: blue; 90%: cyan; 70%: yellow; 50%: orange; 0%: red). The pLDDT score for each key residue is shown.

**Figure S4.**
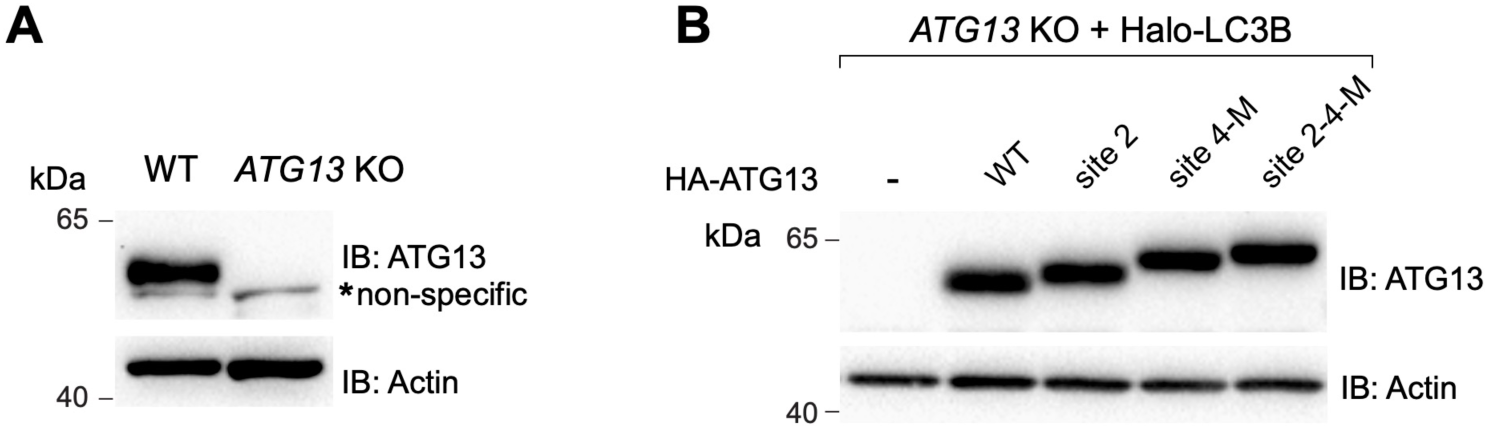
Generation of knockout and rescue cell lines, related to Figure 2. (**A**) Cell lysates from WT and *ATG13* KO HeLa cells were analyzed by immunoblotting with indicated antibodies. (**B**) Cell lysates from *ATG13* KO cells expressing Halo-LC3B without rescue or rescued with indicated versions of HA-ATG13 were immunoblotted with anti-ATG13 and anti-Actin antibodies.

**Figure S5.**
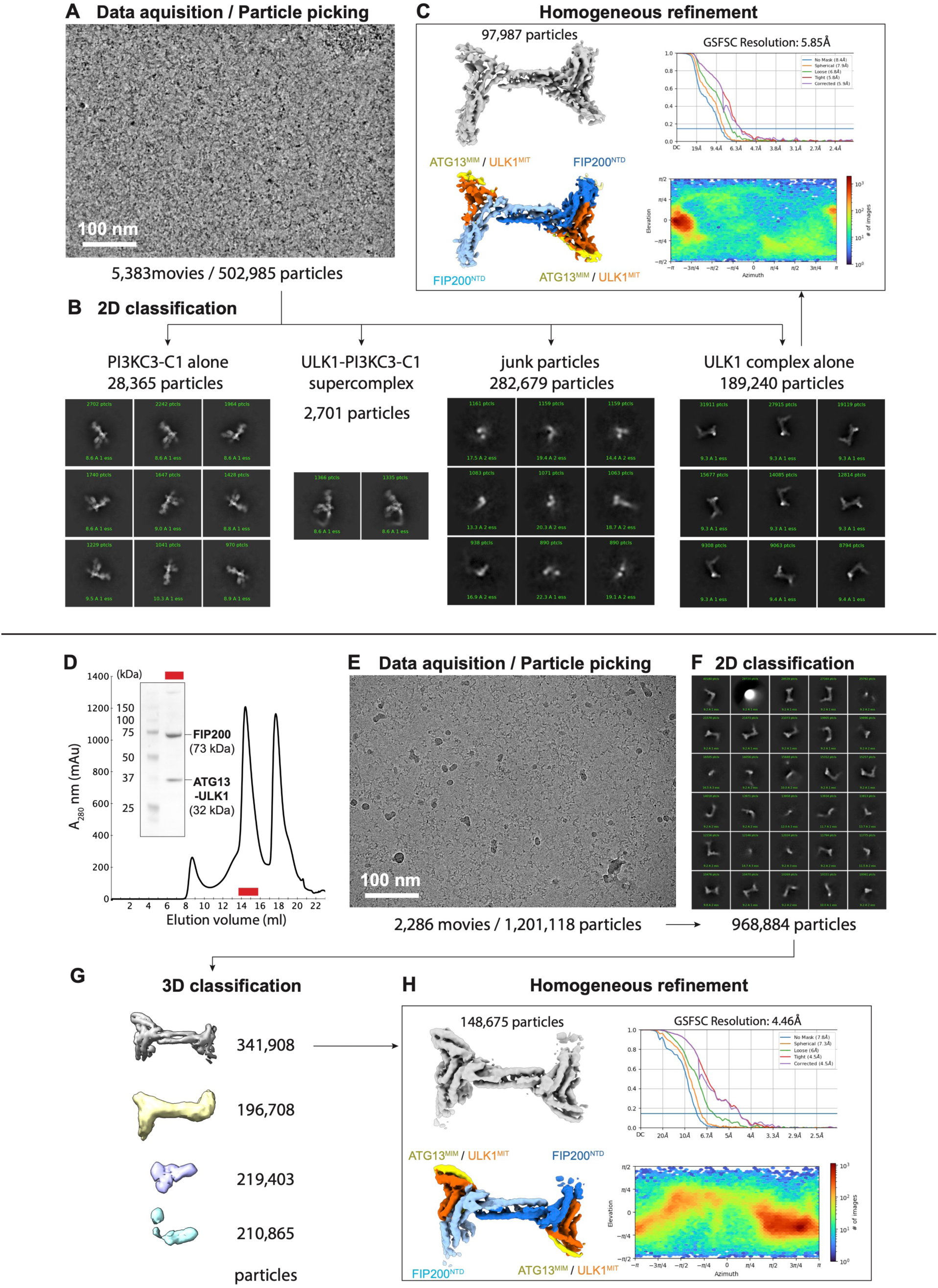
Structural determination of the the ULK1C (2:2:2) core in the PI3KC3-C1 mixture and the ATG13^450–517^truncation mutant, related to Figure 4 and Figure 5. (**A**) A representative cryo-EM micrograph of the ULK1C:PI3KC3-C1 mixture sample. (**B**) Representative 2D class averages of four subpopulations of the dataset. From left: PI3KC3-C1 alone, ULK1C:PI3KC3-C1 supercomplex, junk particles, and ULK1C alone. (**C**) Result of the homogeneous refinement of the ULK1C alone substack. The FSC, local resolution, and angular distribution are shown accordingly. The EM map is contoured at 7σ. (**D**) Size exclusion chromatography (SEC) profile of the ULK1C (2:2:2) core of ATG13^450–517^ truncation mutant. The inset shows an SDS-PAGE of the peak (red bar). (**E**) A representative cryo-EM micrograph of the ULK1C core containing the ATG13^450–517^ truncation mutant. (**F**) Representative 2D class averages. (**G**) Result of the first round of 3D classification. (**H**) Global refinement of the final substack. The FSC, local resolution, and angular distribution are shown accordingly.

**Figure S6.**
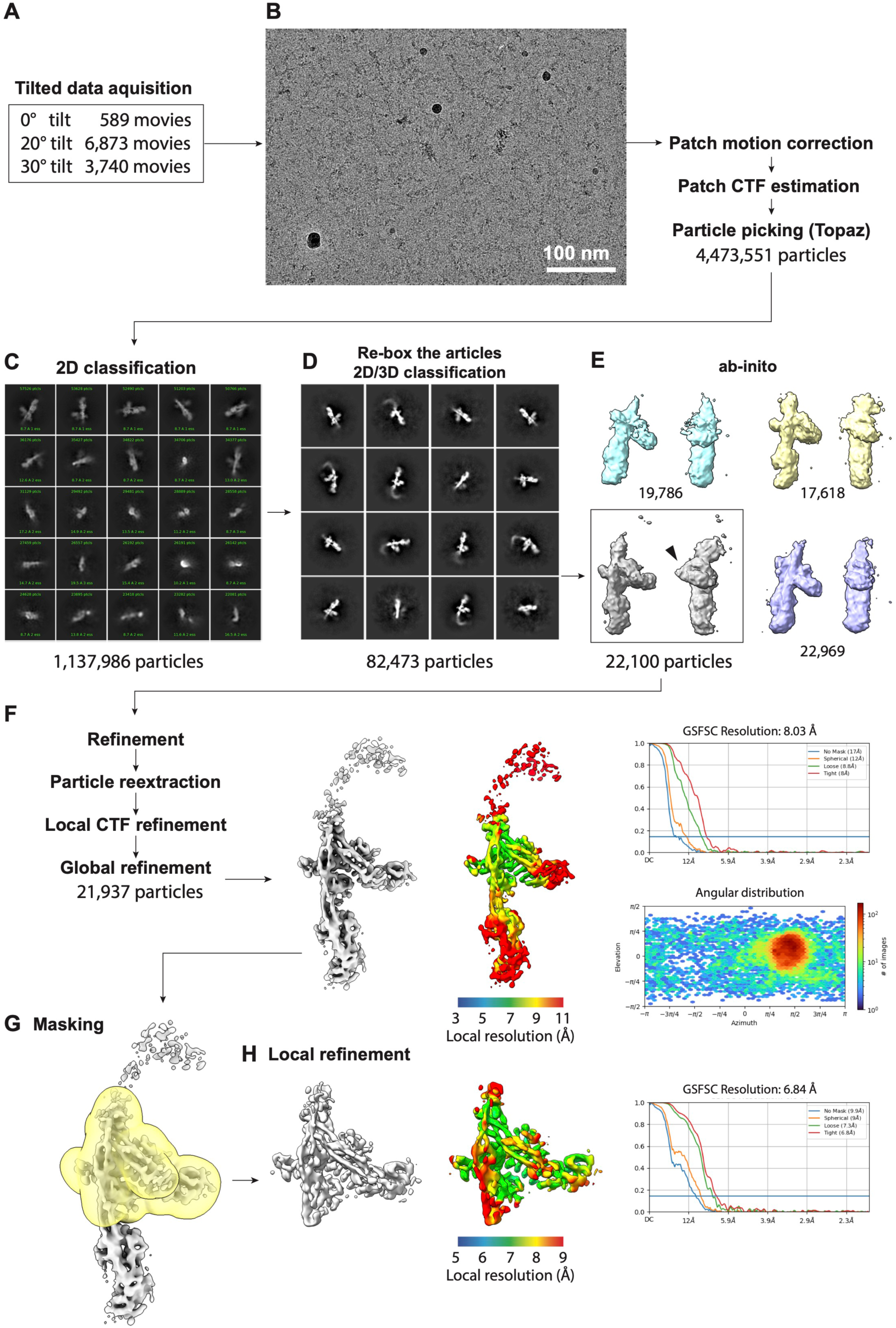
Structural determination of the ULK1C:PI3KC3-C1 supercomplex, related to Figure 4. (**A**) Dataset acquisition. Three data sets were collected at 0°, 20, and 30° tilted specimen stage. The number of movies collected for each angle is shown respectively. (**B**) A representative cryo-EM micrograph is shown on the left. Scale bar: 100 nm. (**C**) Representative 2D class averages of the first round of 2D classification. (**D**) Representative 2D class averages of the intermediate particle stack, re-extracted with a larger box size, and sorted by multiple rounds of 2D and 3D classification. (**E**) The initial model created from the particle stack. The black arrow indicates the EM density of FIP200. (**F**) Result of the homogeneous refinement. The local resolution, FSC, and angular distribution are shown accordingly. The map is displayed at a contour level of 8σ. (**G**) Result of the local refinement. The mask for local refinement, local resolution, FSC, and angular distribution are shown accordingly. The map is displayed at a contour level of 17σ.

**Figure S7.**
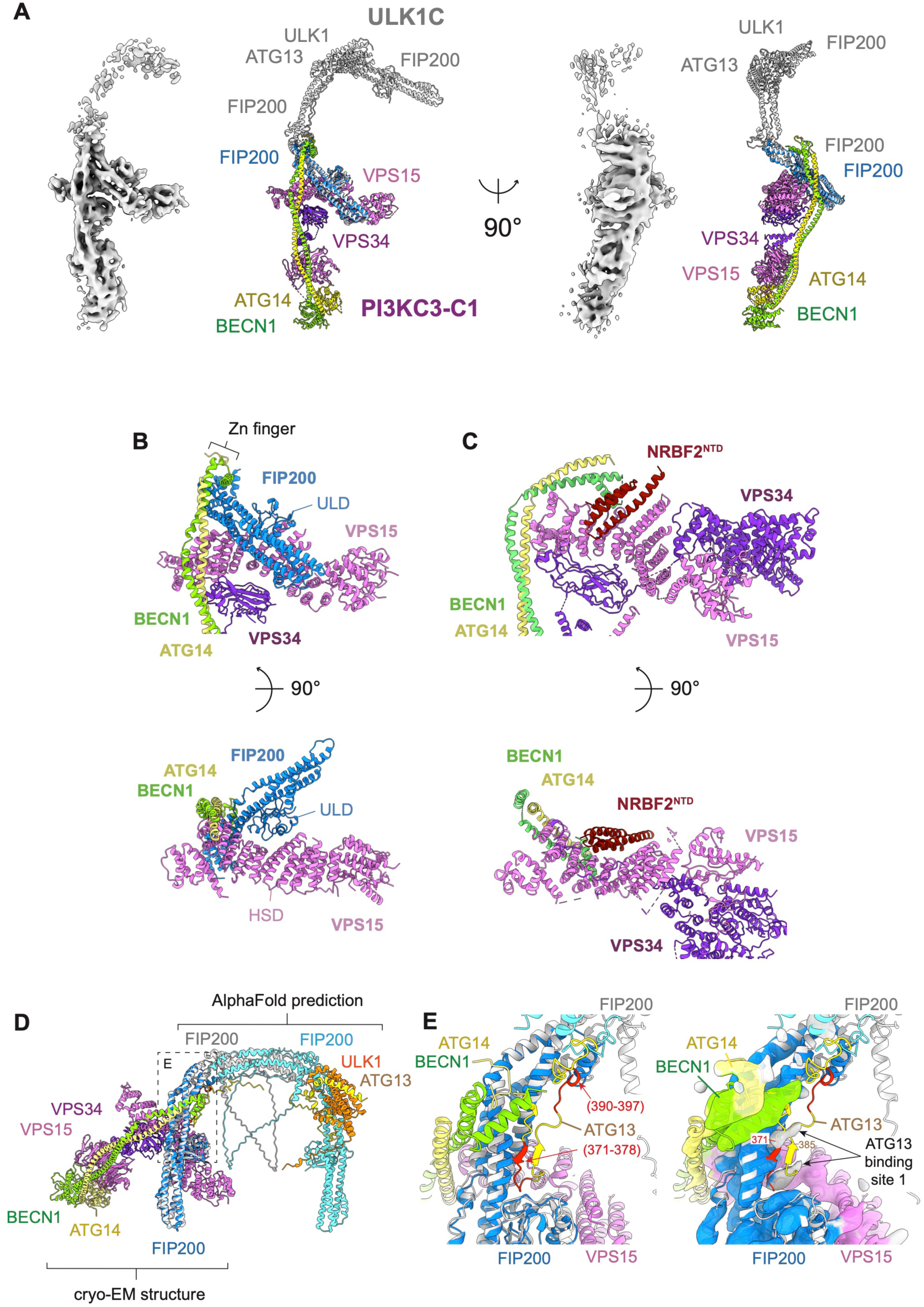
Characterization of the ULK1C:PI3KC3-C1 supercomplex structure, related to Figure 4. (**A**) Overview of the ULK1C:PI3KC3-C1 supercomplex. The cryo-EM structure of the ULK1C core (2:1:1) determined in this study is superposed onto the FIP200 of the supercomplex and colored in gray. **(B** and **C**) Comparison of the VPS15^HSD^ binding sites in binding with FIP200 (blue) and NRBF2 (red) (*51*). (**D**) Superposition of the ULK1C core (2:1:1) prediction model onto the supercomplex cryo-EM structure. The FIP200 molecule of the prediction model (gray) is superposed to the FIP200 molecule of the supercomplex (blue). (**E**) Close up view of the FIP200 site 1 and 2. The FIP200 interacting regions of ATG13 (371-378, 390-397), identified with HDX in the previous study (*52*) are colored red. (Right) The same view with showing the local refinement EM map at 22.5σ. The two extra EM maps observed at site 1 correspond to the predicted ATG13 region 371-385.

**Figure S8.**
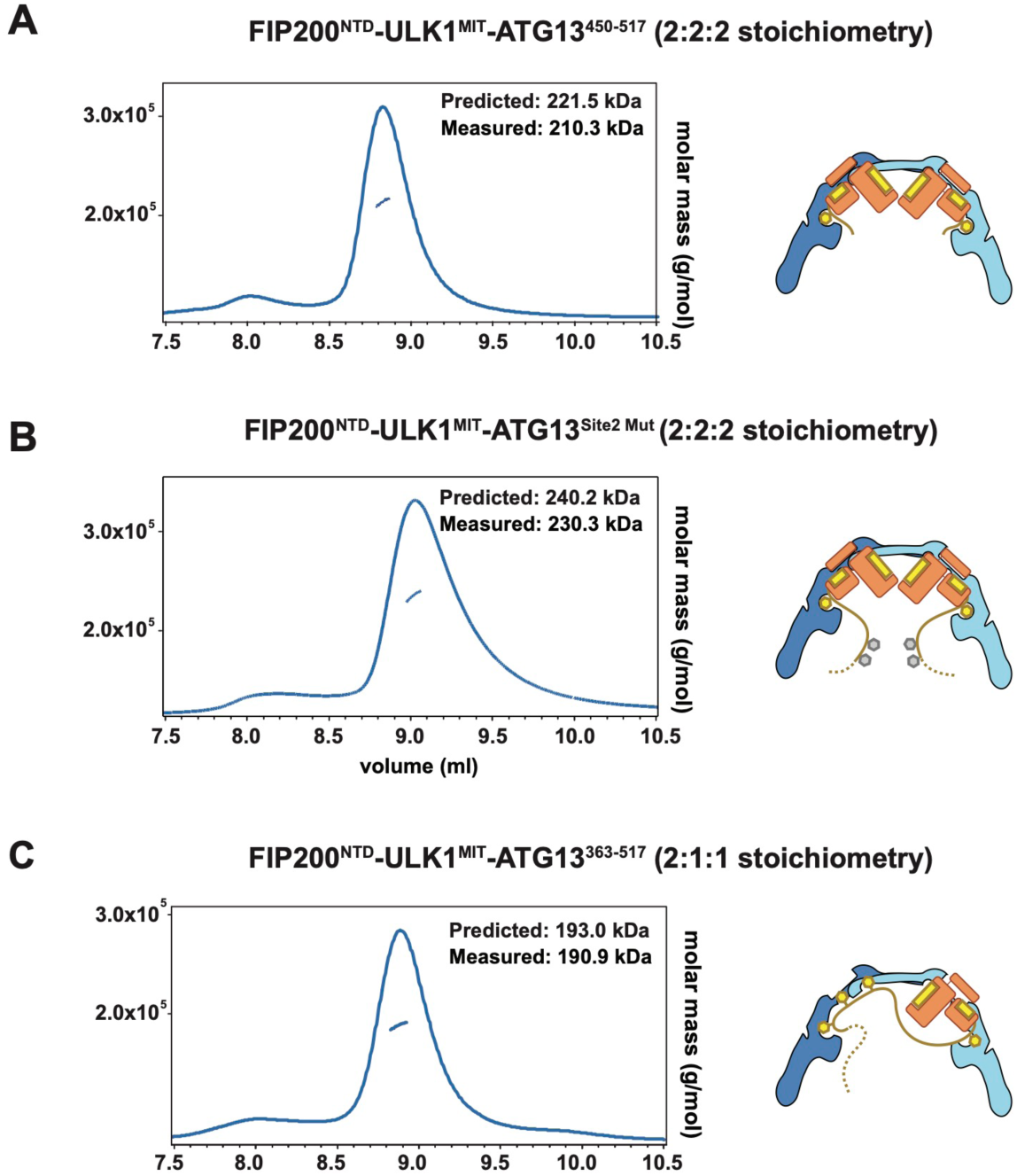
ATG13^IDR^ regulates the 2:2:2 complex formation of the ULK1C, related to. **Figure 5**. (A) SEC-MALS result of the ULK1C core containing truncated ATG13 variant (ATG13^450–517^), (B) SEC-MALS result of the ULK1C core containing site 2 mutated ATG13 variant (ATG13^363– 517^ with mutations F394D/F397D/E403K), (**C**) SEC-MALS result of the ULK1C core containing wild type ATG13 fragment (ATG13^363–517^). The predicted and measured molecular weight for each measurement are shown. The schematic drawing illustrates the predicted assembly of each condition.

**Table S1:** Cryo-EM data collection and refinement statistics, related to the STAR Methods.

|  | ULK1C core<br>(2:1:1)<br>stoichiometry | ULK1C:PI3KC3-C1<br>supercomplex | ULK1C core<br>(2:2:2) stoichiometry<br>in PI3KC3-C1 mixture | ULK1C core<br>(2:2:2) stoichiometry<br>ATG13<sup>450-517</sup> mutant |
| --- | --- | --- | --- | --- |
|  | EMD-40658<br>8SOI | EMD-45297<br>9C82 | EMD-40715<br>8SQZ | EMD-40735<br>8SRM |
| **Data collection and processing** |  |  |  |  |
| Magnification | 81,000 | 81,000 | 81,000 | 36,000 |
| Voltage (kV) | 300 | 300 | 300 | 200 |
| Electron exposure (e<sup>-</sup>/Å<sup>2</sup>) | 50 | 50 | 50 | 50 |
| Defocus range (μm) | -0.8 to -2.0 | -0.8 to -2.0 | -0.8 to -2.0 | -0.8 to -2.0 |
| Physical pixel size (Å) | 1.05 | 1.05 | 1.05 | 1.115 |
| Symmetry imposed | C1 | C1 | C1 | C1 |
| Images | 8,468 | 589 (0° tilt)<br>6,973 (20° tilt)<br>3,740 (30° tilt) | 5,383 | 2,286 |
| Initial particle images (no.) | 2,156,516 | 4,473,551 | 502,985 | 968,884 |
| Final particle images (no.) | 267,719 | 21,937 | 97,987 | 149,409 |
| Map resolution, Global (Å) | 4.2 | 8.03 | 5.28 | 4.46 |
| Map resolution, Best local (Å) | 3.35 | 6.84 | n/a | n/a |
| FSC threshold | 0.143 | 0.143 | 0.143 | 0.143 |
| **Refinement** |  |  |  |  |
| Map sharpening B factor (Å<sup>2</sup>) | 143.2 | 752.0 | 431.6 | 155.4 |
| Model composition |  |  |  |  |
| Non-hydrogen atoms | 10,243 | 11,172 | 7,438 | 5,773 |
| Protein residues | 1,276 | 2,254 | 1,498 | 1,163 |
| Ligands | 0 | 0 | 0 | 0 |
| B factors (Å<sup>2</sup>) |  |  |  |  |
| Protein | 126.43 | 1099.03 | 319.79 | 208.69 |
| Ligands | n/a | n/a | n/a | n/a |
| R.m.s. deviations |  |  |  |  |
| Bond lengths (Å) | 0.003 | 0.002 | 0.004 | 0.002 |
| Bond angles (°) | 0.598 | 0.607 | 0.633 | 0.405 |
| Validation |  |  |  |  |
| MolProbity score | 1.7 | 2.48 | 1.95 | 1.51 |
| Clashscore | 9.52 | 30.51 | 8.07 | 4.44 |
| Poor rotamers (%) | 1.5 | n/a | n/a | n/a |
| Ramachandran plot |  |  |  |  |
| Favored (%) | 96.75 | 91.17 | 91.41 | 95.87 |
| Allowed (%) | 3.17 | 8.83 | 8.59 | 4.13 |
| Disallowed(%) | 0.008 | 0 | 0 | 0 |

**Table S2:** Theoretical and measured mass in mass photometry analysis, related to Figure 6. FIP200n, ATG13c, and ULK1c represent FIP200 (1-640), ATG13 (363-517), and ULK1 (836-1059), respectively.

| Measurement | Sample | Complex | Expected (kDa) | Measured (kDa) | % |
| --- | --- | --- | --- | --- | --- |
| i) | FIP200n | FIP200n monomer | 80 | 85 | 5 |
|  |  | FIP200n dimer | 160 | 166 | 95 |
| ii) | MBP-ATG13c-ULK1c | ATG13-ULK1 monomer | 85 | 87 | 95 |
|  |  | ATG13-ULK1 dimer | 170 | 174 | 5 |
| iii) | PI3KC3-C1 | PI3KC3-C1 | 367 | 377 | 100 |
| iv) | FIP200n +<br>MBP-ATG13(450-517)-<br>ULKc | ATG13-ULK1 monomer | 77 | 81 | 40 |
|  |  | FIP200n dimer | 160 | 157 | 15 |
|  |  | ULK1C 2:1:1 complex | 237 | 247 | 20 |
|  |  | ULK1C 2:2:2 complex | 314 | 318 | 25 |
| v) | FIP200n +<br>MBP-ATG13c-ULK1c<br>Crosslinked at 10 nM | ATG13-ULK1 monomer | 85 | 89 | 56 |
|  |  | FIP200n dimer | 160 | 170 | 7 |
|  |  | ULK1C 2:1:1 complex | 245 | 252 | 33 |
|  |  | ULK1C 2:2:2 complex | 330 | 338 | 4 |
| vi) | FIP200n +<br>MBP-ATG13c-ULK1c +<br>PI3KC3-C1,<br>Crosslinked at 10 nM | ATG13-ULK1 monomer | 85 | 88 | 43 |
|  |  | FIP200n dimer | 160 | 171 | 5 |
|  |  | ULK1C 2:1:1 complex | 245 | 260 | 30 |
|  |  | ULK1C 2:2:2 complex | 330 | 358 | 5 |
|  |  | PI3KC3-C1 | 367 | 387 | 16 |
| vii) | FIP200n +<br>MBP-ATG13c-ULK1c,<br>Crosslinked at 200nM | ATG13-ULK1 monomer | 85 | 89 | 36 |
|  |  | FIP200n dimer | 160 | 170 | 12 |
|  |  | ULK1C 2:1:1 complex | 245 | 254 | 35 |
|  |  | ULK1C 2:2:2 complex | 330 | 338 | 17 |
| viii) | FIP200n +<br>MBP-ATG13c-ULK1c +<br>PI3K-C1,<br>Crosslinked at 200nM | ATG13-ULK1 monomer | 85 | 89 | 39 |
|  |  | FIP200n dimer | 160 | 177 | 8 |
|  |  | ULK1C 2:1:1 complex | 245 | 249 | 29 |
|  |  | ULK1C 2:2:2 complex | 330 | 330 | 13 |
|  |  | PI3KC3-C1-C1 | 367 | 380 | 11 |
| ix) | FIP200n +<br>MBP-ATG13c-ULK1c,<br>Crosslinked at 1000nM | ATG13-ULK1 monomer | 85 | 91 | 31 |
|  |  | FIP200n dimer | 160 | 164 | 10 |
|  |  | ULK1C 2:1:1 complex | 245 | 254 | 25 |
|  |  | ULK1C 2:2:2 complex | 330 | 335 | 33 |
| x) | FIP200n +<br>MBP-ATG13c-ULK1c +<br>PI3KC3-C1,<br>Crosslinked at 1000nM | ATG13-ULK1 monomer | 85 | 93 | 23 |
|  |  | FIP200n dimer | 160 | 162 | 13 |
|  |  | ULK1C 2:1:1 complex | 245 | 257 | 18 |
|  |  | ULK1C 2:2:2 complex | 330 | 346 | 23 |
|  |  | PI3KC3-C1 | 367 | 383 | 25 |

